# Dual S100A1 and ARC gene therapy as a treatment for DMD cardiomyopathy

**DOI:** 10.1101/2025.08.23.671924

**Authors:** David W. Hammers, Cora C. Hart, Young il Lee, Margaret M. Sleeper, H. Lee Sweeney

## Abstract

Duchenne muscular dystrophy (DMD) is a lethal pediatric striated muscle disease caused by loss of dystrophin for which there is no cure. Cardiomyopathy is the leading cause of death amongst individuals with DMD, and effective therapeutics to treat DMD cardiomyopathy are a major unmet clinical need. This work investigated adeno-associated viral (AAV) gene therapy approaches to treat DMD cardiomyopathy by overexpression of the calcium binding proteins S100A1 and apoptosis repressor with caspase recruitment domains (ARC). Using the severe D2*.mdx* mouse model of DMD, we identified that S100A1 gene therapy improves the diastolic dysfunction associated with DMD cardiomyopathy, whereas ARC gene therapy prolongs survival. The combination of both S100A1 and ARC in a single bicistronic vector improves the long-term cardiac outcome of D2.*mdx* mice, development of heart failure caused by micro-dystrophin expression, and exhibits safety via intracoronary delivery in a canine model of DMD. Furthermore, S100A1-ARC gene therapy provides functional benefits when expressed in D2.*mdx* skeletal muscle. Together, these findings indicate that S100A1-ARC gene therapy represents an effective treatment for DMD cardiomyopathy and may be effective in treating other forms of cardiomyopathy and muscle pathologies.

**SIGNIFICANCE STATEMENT:** Cardiomyopathy is the leading cause of death amongst individuals with Duchenne muscular dystrophy (DMD). Effective therapeutics to treat DMD cardiomyopathy represent a major unmet clinical need. This work identifies the dual gene therapy approach of S100A1 and ARC as an effective treatment that improves long-term cardiac function and life-expectancy in severe mouse model of DMD. Intracoronary delivery of this AAV-based gene therapy also exhibits safety and evidence of efficacy in dystrophic canines. Furthermore, functional benefits in skeletal muscle are also incurred via S100A1-ARC expression in striated muscle. These findings indicate that S100A1-ARC therapy is an effective treatment for DMD cardiomyopathy whose benefits may be applicable for other forms of cardiac and muscle disease.

## INTRODUCTION

Duchenne muscular dystrophy (DMD) is a lethal X-linked pediatric muscle disease caused by *DMD* mutations that result in loss of dystrophin, a protein required for the stabilization of muscle during contractile activity^1, 2^. Respiratory care advancements for DMD has increased the life expectancy for affected individuals^3, 4^, and, as a consequence, cardiomyopathy has emerged as the leading cause of DMD mortality^5^. Currently, effective therapeutics to combat dystrophic cardiomyopathy are a major unmet clinical need for individuals with DMD.

Prominent aspects of DMD cardiomyopathy include electrical conduction abnormalities and arrhythmic events that can be detected early in life^4, 6–8^ and the gradual progression to dilated cardiomyopathy (DCM) in the second to third decade of life^9^. A feature of DMD cardiomyopathy receiving increased recognition is a prolonged phase of left ventricular (LV) restriction and pronounced diastolic dysfunction that precedes overt decline of systolic function^10–13^. While stroke volume (SV) deficits and LV wall movement abnormalities have been reported in younger DMD boys as early as 1979^8^, advanced echocardiography and cardiac magnetic resonance (CMR) techniques yield data clearly depicting this restrictive phenotype in DMD patients prior to EF decline^11, 13–15^. Su *et al.*^11^ brought specific attention to this LV restriction in DMD, coined as “tonic contraction”, and the recent work of Starnes *et al.*^12^ demonstrates clear impairments of DMD LV filling using CMR. In agreement with these clinical findings, diastolic impairments are also evident in mouse and dog models of DMD^16–19^ (**Figure 1** and **Table S3**).

**Figure 1.**
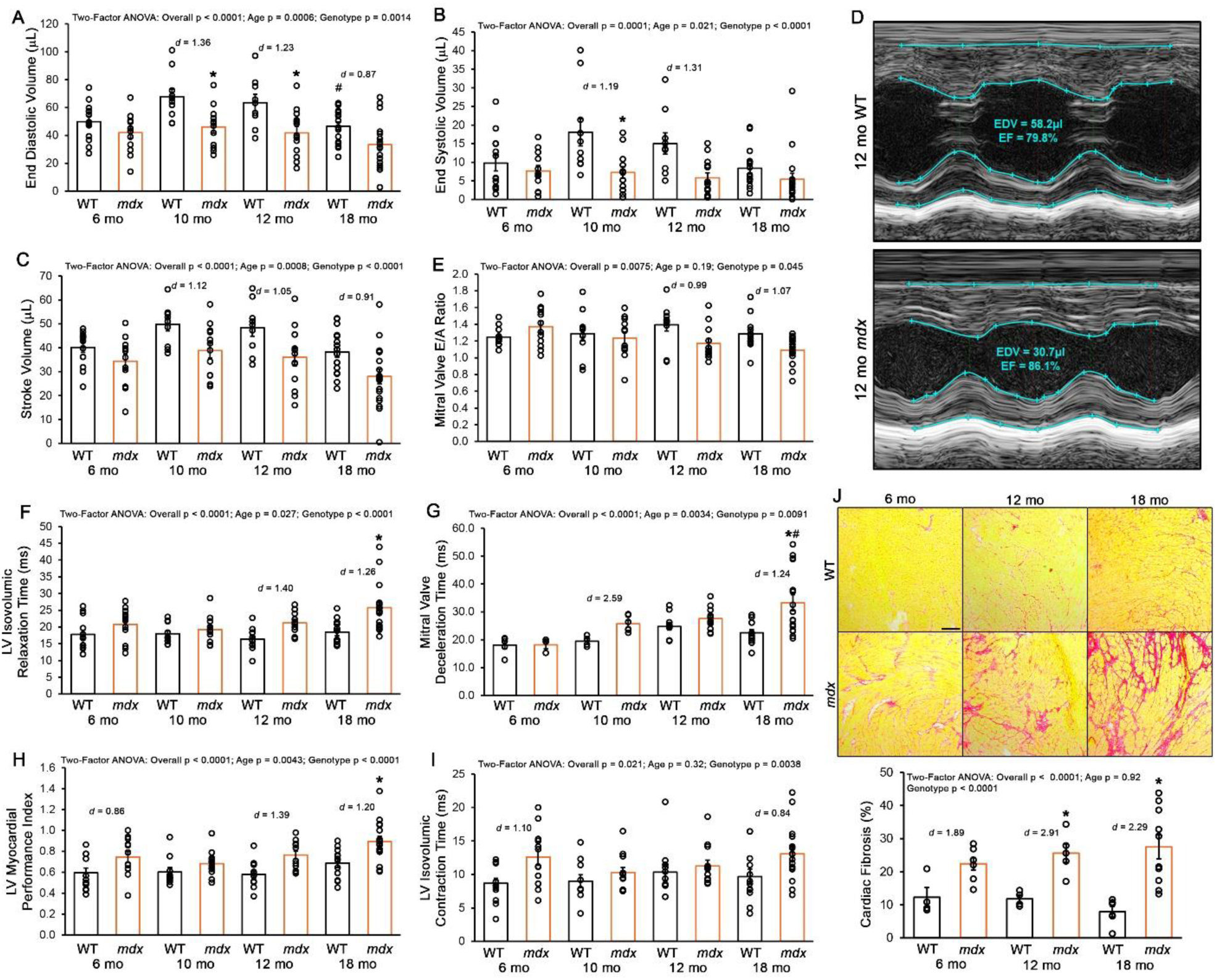
Progressive diastolic dysfunction and LV restriction in D2.*mdx* mice. Echocardiography was performed on D2.WT (WT; n = 9-17) and D2.*mdx* (*mdx*; n = 12-18) mice at the ages of 6, 10, 12, and 18 months of age (mo). Measures of left ventricular (LV) (**A**) end diastolic volume, (**B**) end systolic volume, and (**C**) stroke volume were determined from (**D**) short-axis M-mode images. Pulsed wave Doppler was used to determine (**E**) mitral valve (MV) E/A Ratios, (**F**) LV isovolumic relaxation time, (**G**) MV deceleration time, (**H**) LV myocardial performance index, and (**I**) LV isovolumic contraction time. (**J**) Cardiac fibrosis was visualized and quantified using picrosirius red staining of heart sections from 6, 12, and 18 mo mice of each genotype (scale bar represents 100 µm). Data are displayed as mean±SEM with individual values indicated by open circles and were analyzed using two-factor ANOVA (age and genotype effects) followed by Tukey post-hoc tests (α = 0.05). *p < 0.05 vs. age-matched WT values; #p < 0.05 vs. genotype-matched values from the previous age group; Large effect sizes between age-matched genotypes are also indicated.

Mechanisms contributing to the development of DMD cardiomyopathy largely converge on the central hypothesis that loss of dystrophin causes impaired cardiomyocyte calcium handling^20–22^. This model entails persistently high intracellular calcium concentrations caused by sarcolemma tears, dysfunctional cation channels, sarcoplasmic reticulum (SR) leakage, and/or insufficient removal of calcium from the cytosol result in a myriad of detrimental consequences, including (but not limited to) impaired relaxation, arrhythmogenic events, protease activation, mitochondrial dysfunction, and cell death^17, 22–25^. To address this pathogenic calcium dysregulation in dystrophic cardiomyocytes, several strategies have been employed to restore calcium homeostasis, including increasing SERCA levels/activity^26–28^, addressing SR calcium leak through ryanodine receptors (RyRs)^29, 30^, blocking calcium-permeable cation channels^31, 32^, and preventing sarcolemmal tears with membrane sealants^33, 34^.

The purpose of the current study is to investigate the impact of S100A1, a calcium binding protein that exerts several benefits in context of calcium dysregulation during heart failure (HF)^35^, and apoptosis repressor with caspase recruitment domain (ARC), an anti-apoptotic protein that improves cardiomyocyte survival during myocardial infarction^36–38^, as novel gene therapies for DMD cardiomyopathy. Herein, we provide the time course of D2.*mdx* cardiac progression to a restrictive phenotype with severe diastolic dysfunction. This phenotype is effectively prevented by adeno-associated virus (AAV)-mediated cardiac S100A1 overexpression. AAV-mediated cardiac ARC overexpression, on the other hand, does not largely affect LV restriction, but does prolong the survival of D2.*mdx* mice. Furthermore, a cardiac bicistronic AAV containing both S100A1 and ARC improves long-term cardiac outcomes of D2.*mdx* mice, prevents HF associated with high levels of micro-dystrophin, and exhibits safety and evidence of cardiac improvements in the GRMD canine model of DMD. Together, these data indicate that the combinatorial delivery of S100A1 and ARC utilizing a bicistronic AAV approach represents a novel gene therapy strategy for the treatment of DMD cardiomyopathy.

## METHODS

### Animals

All animal studies were approved and conducted in accordance with the University of Florida IACUC. This study used male wild-type (D2.WT; Jax# 000671) and *mdx* (D2.*mdx*; Jax# 013141) mice on the DBA/2J genetic background from colonies originally obtained from Jackson Laboratory^39, 40^. Mice were housed 1-5 mice per cage, randomly assigned into groups, provided *ad libitum* access to food (NIH-31 Open formulation diet; Envigo #7917), water, and enrichment, and maintained on a 12-hour light/dark system. GRMD canines were housed and cared for as previously described^17, 41^. Mouse treatments entailed IV delivery of vector via tail-vein injections, as previously described^16^. Canine intracoronary vector delivery methods are described Supplemental Methods.

### AAV vector production

Codon-optimized murine S100A1 and ARC open reading frames were synthesized by Genscript (Piscataway, NJ) and cloned into a pAAV shuttle plasmid containing the cardiac-specific cTnT promoter^42^ and a minimized synthetic polyadenylation signal sequence^43^. The bicistronic construct contained native canine S100A1, an internal ribosomal entry site (IRES)^44^, and codon-optimized canine ARC sequences, and were driven by either the cTnT or CK8e^45^ promoters. AAV viral vector packaging was performed using the triple-transfection method, as previously described^46, 47^.

### Echocardiography and electrocardiograms

Mouse electrocardiograms and transthoracic echocardiograms were performed as previously described^16^ using the Vevo 3100 pre-clinical imaging system (Fujifilm Visualsonics). Details of these procedures are found in the Supplemental Methods. Echocardiography measurements in canines were performed, as described previously^17^, using a Philips CX-50 system (Philips) and an 8- to 3-MHz transducer.

### Ex vivo muscle function

Maximal tetanic tension assessments of the extensor digitorum longus (EDL) and soleus muscles were evaluated as previously described^40, 41^ by the University of Florida Physiological Assessment Core. Following experimental procedures, muscles were weighed, frozen in OCT or snap-frozen, and stored at -80 C until further use.

### Immunoblotting

Immunoblotting was performed as previously described^39, 48^ using the following primary antibodies: anti-S100A1 (Abcam #11428), anti-ARC/NOL3 (Proteintech# 10846-2-AP), anti-dystrophin [DSHB# MANEX1011B(1C7)], and anti-GFP (Abcam# 13970). Enhanced chemiluminescent signals were acquired and quantified using the Licor C-Digit system, with signal intensity normalized to sample protein loading, as visualized using Ponceau Red staining.

*Gene Expression and Vector Genome Quantification* Gene expression analysis was conducted as previously described^39, 48^ using the following mouse-specific primers: *Nppb* (forward) 5’-TTT GGG CTG TAA CGC ACT GA-3’ and (reverse) 5’-CAC TTC AAA GGT GGT CCC AGA-3’; *Myh6* (forward) 5’-AAC CTG TCC AAG TTC CGC A-3’ and (reverse) 5’-ATT CCT CGT CGT GCA TCT TCT TG-3’; *Myh7* (forward) 5’-TTG CTA CCC TCA GGT GGC T -3’ and (reverse) 5’-CCT TCT CAG ACT TCC GC AGG -3’; *Rpl32* (forward) 5’-GGG TGC GGA GAA GGT TCA A-3’ and (reverse) 5’-GCA ATC TCA GCA CAG TAA GAT TTG T-3’. Relative gene expression quantification was performed using the ΔΔCt method with *Rpl32* as the normalization gene.

Vector genome content quantification was performed as previously described^16^. Primers used during this assay include those for AAV2 ITR (recognizes vector genomes; Forward: 5’-GGA ACC CCT AGT GAT GGA GTT-3’; Reverse: 5’-CGG CCT CAG TGA GCG A-3’) and a genomic DNA region in the *Rpl32* locus of murine chromosome 6 (recognizes diploid genomes; Forward: 5’-GAG AAG GTT CAA GGG CCA GAT -3’; Reverse: 5’-AGC TCC TTG ACA TTG TGG ACC-3’). Canine diploid genomes were quantified using primers specific for the genomic DNA regions in the *FUCA1* locus of canine chromosome 2 (Forward: 5’-CGC GCT TCT TCC ACC CCGACA CCT-3’; Reverse: 5’-CCC CCG CCC CGA GCA GAC G-3’). Vector genome expression was quantified normalized to diploid genome expression using the ΔΔCT method.

### Histology

Hematoxylin & Eosin (H&E) and picrosirius red (PSR) staining was performed as previously described^39, 40^. Slides were visualized with a Leica DMR microscope, and images were acquired using a Leica DFC310FX camera interfaced with Leica LAS X software. Images were processed and analyzed by investigators blinded to study groups using ImageJ software.

### Statistical analysis

Statistical analysis was performed using unpaired, two-tailed Welch’s T-test (α = 0.05), ANOVA (one-factor, two-factor, or repeated measures) followed by Tukey HSD post-hoc tests (α = 0.05), and Kaplan-Meier estimator analyses (α = 0.05), where appropriate. A P value less than 0.05 was considered significant. Data are displayed as mean ± SEM with individual values indicated or survival curves. Effect size between specific groups is indicated by Cohen’s *d* (*d*).

## RESULTS

### Progressive diastolic dysfunction and LV restriction in D2.mdx mice

The mild progressing course of disease in traditional *mdx* mouse models of DMD requires advanced aging or use of stress-inducing stimuli to elicit functional deficits^49, 50^, thereby has limited mechanistic insights into and therapeutic development for DMD cardiomyopathy. We have previously described and demonstrated the suitability of the D2.*mdx* mouse to model and evaluate translational therapeutics for the severe pathological features of DMD skeletal muscle^16, 39, 40, 48^. To gain a better insight of this murine model’s utility for understanding and testing therapeutic interventions for DMD cardiomyopathy, we performed a comprehensive evaluation of the development of cardiomyopathy in D2.*mdx* mice using electrocardiography (ECG), echocardiography, and histology.

Electrical abnormalities detected using ECG are typically the first clinical symptoms of DMD cardiomyopathy^6–8^, therefore we analyzed ECGs from wild-type (WT) and *mdx* mice of the DBA/2J genetic background from 2 to 12 months of age (mo). Similar to observations in younger boys with DMD, ECG abnormalities are evident in D2.*mdx* as early as 2 mo (**Table S1**). This includes significant genotype effects for prolonged QT, JT, and T_peak_ to T_end_ intervals, as well as increased T wave amplitude, which are indicative of impaired LV repolarization. These data confirm that, like DMD boys, D2.*mdx* mice exhibit electrical abnormalities early in life.

Diastolic dysfunction and restrictive LV filling are becoming more frequent aspects reported of DMD cardiomyopathy prior to onset of systolic deficits and DCM development^10–13, 15^. To analyze the emergence of these features in D2.*mdx* mice, we performed echocardiography at 6, 10, 12, and 18 mo. Compared to their WT counterparts, D2.*mdx* mice exhibit progressive declines in end diastolic volume (EDV), end systolic volume (ESV), and stroke volume (SV) beginning at 10 mo, with significant age and genotype effects (**Figure 1A-D**). Four-chamber pulse-wave Doppler measurements reveal significant *mdx* associated reductions in mitral valve (MV) E/A ratio (**Figure 1E**), a key indicator of diastolic dysfunction, as well as significant age and genotype associated increases LV isovolumic relaxation time (IVRT; **Figure 1F**), MV deceleration time (**Figure 1G**), and LV myocardial performance index (MPI; **Figure 1H**), also measures of impaired LV relaxation and diastolic filling. LV isovolumic contraction time (IVCT) was also significantly increased in *mdx* mice (**Figure 1I**), suggesting defects in LV contractility, pressure development, and/or aortic valve opening may be aspects of cardiomyopathy in the D2.*mdx* mouse. These measures of impaired *mdx* diastolic function are not due to faster heart rates than WT counterparts (**Figure S1A**). While there were no significant differences in ejection fraction (EF) across groups, individual *mdx* mice in the 18 mo age group exhibit substantially lower EFs than the rest of the group (**Figure S1B**). Although evidence of LV wall thinning at diastole are not observed to confirm progression to DCM in those mice (**Figure S1C-D**), these instances of reduced EF may indicate that systolic failure begins around 18 mo in this model. While some features of LV diastolic dysfunction and restriction may be interpreted as development of cardiac hypertrophy, we identified no evidence of *mdx* associated cardiac enlargement at the organ or cardiomyocyte levels in 12 mo mice (**Table S2**). Furthermore, we identified many of these LV restriction phenotypes in historical data from GRMD dogs, an established large animal model to study DMD cardiomyopathy^17, 51, 52^. This analysis revealed affected GRMD dogs exhibit reduced LV end diastolic parameters, SVs, and MV E/A ratios compared to normal and carrier canines (**Table S3**). These data indicate that diastolic dysfunction with LV restriction is a translational feature of DMD cardiomyopathy that is prominent in the D2.*mdx* mouse model of DMD.

Consistent with the severe histopathology exhibited by D2.*mdx* skeletal muscle^40^, the hearts of these mice also show increased fibrosis and tissue pathology, as demonstrated by PSR staining (**Figure 1J**). This cardiac fibrosis is consistent with features found in DMD boys and most preclinical models of DMD^17, 53–56^ and is a potential mechanism contributing to LV restriction, as it may serve as a physical barrier to proper cardiomyocyte elongation during diastole. To test this hypothesis, we acutely treated 10 mo *mdx* mice exhibiting reduced EDVs with the cardiac myosin inhibitor mavacamten^57^ (also known as MYK-461; 2.5 mg/kg; experiment depicted in **Figure S2A**) to determine if **a**) manipulating (intrinsic) cardiomyocyte contractile activity of restricted *mdx* mice can abrogate the reduced EDV or **b**) extrinsic factors, such as fibrosis, impede LV diastolic opening. As anticipated by its mechanism of action, mavacamten decreased fractional shortening by 26% 3 hours following oral administration, and this effect was completely reversed following a 3 day washout period (**Figure S2B-C**). Mavacamten also acutely increased EDV by 68%, which was also reversed by a 3 day washout period. The acutely reversible nature of LV restriction by mavacamten indicates that this phenomenon is intrinsic to cardiomyocytes, not extracellular factors such as fibrosis. Therefore, intracellular mechanisms, such as the rise in resting calcium levels we have previously observed^16^, are responsible for the LV diastolic dysfunction and restrictive phenotype found in D2.*mdx* hearts, which potentially represents a targetable feature to treat DMD cardiomyopathy.

### S100A1 treatment alleviates LV restriction and ARC prolongs life span in D2.mdx mice

Multiple gene therapies have been designed to address calcium handling issues in DMD cardiomyopathy by increasing SERCA activity^26–28^. While effective in mitigating elevated cytosolic calcium concentrations, these strategies also increase ATP demands in already energetically challenged dystrophic cardiomyocytes^20^. As an alternative approach to address this calcium dysregulation with gene therapy, we sought to individually test the efficacy of overexpressing either S100A1, a pleiotropic calcium-binding protein that exerts several benefits in failing cardiomyocytes^35^, or ARC, a protein that represses calcium dysregulation’s ultimate downstream effect of cell death^36, 37, 58^.

To confirm that S100A1 and/or ARC are not naturally upregulated during dystrophic cardiomyopathy, immunoblotting for these proteins in 12 mo *mdx* hearts reveals trends of downregulation relative to age-matched WT levels (**Figure S3**). To test the hypothesis that overexpression of each of these proteins would benefit dystrophic hearts, we developed individual self-complementary AAV vectors containing codon-optimized murine ARC and S100A1 driven by the cardiac troponin T (cTnT; *Tnnt2* gene) promoter^42^ and packaged in AAVrh10 serotype vector^46, 59^. These vectors were administered to 1 mo male *mdx* mice via tail vein injections using a dose of 6.33×10^13^ gc/kg (experiment depicted in **Figure 2A**), with echocardiography performed at 12 mo and 18-22 mo (endpoint determined when S100A1 treatment mice reached the point of mean survival). Cardiac AAV genome content was comparable between ARC and S100A1 treatment groups (**Figure 2B**). Both transgenes extend mean survival time beyond the course of untreated *mdx* mice, however, this effect was significant with ARC overexpression(**Figure 2D**). While both S100A1 and ARC improved LV histopathology compared to untreated hearts, ARC treatment led to better cardiomyocyte organization and less signs of infiltration than S100A1 treatment, which agrees with this extension of lifespan (**Figure 2D**).

**Figure 2.**
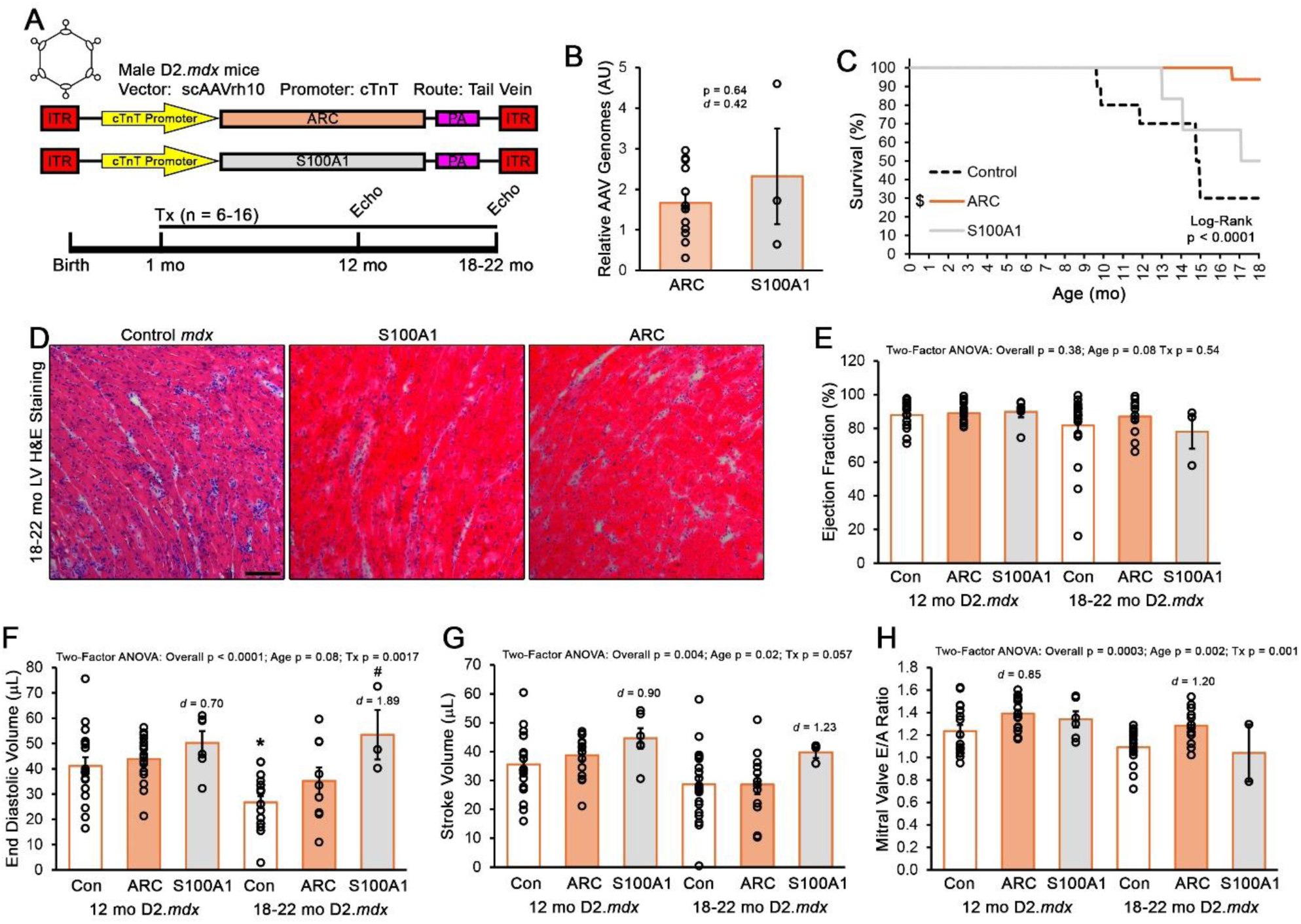
Therapeutic effect of S100A1 and ARC in D2.*mdx* mouse hearts. (**A**) Male D2.*mdx* received systemic treatment with 6.33×10^13^ gc/kg scAAVrh10.cTnT vector containing the coding sequence for either S100A1 (n = 6) or ARC (n = 16) at 1 month of age (mo) and underwent echocardiography at 12 and 18-22 mo time points. Untreated control D2.*mdx* mice (n = 10) were also included in this cohort for survival analysis, whereas additional controls are included in echocardiography measurements. (**B**) Vector transduction was assessed in the hearts of treated mice at endpoint. (**C**) Mouse survival of this study is depicted in a Kaplan-Meier plot. (**D**) Representative images of LV histopathology are shown, as visualized by hematoxylin & eosin (H&E) staining (scale bar represents 100 µm). Echocardiography determined values for (**E**) ejection fraction, (**F**) end diastolic volume, (**G**) stroke volume, and (**H**) mitral valve E/A ratio for 12 mo and 18-22 mo timepoints. Data are displayed as mean±SEM with individual values indicated by open circles and were analyzed using two-factor ANOVA (age and genotype effects) followed by Tukey post-hoc tests (α = 0.05), unless otherwise indicated. *p < 0.05 vs. treatment-matched values from previous timepoint; #p < 0.05 vs. age-matched control values; Large effect sizes from age-matched control groups are also indicated.

While echocardiography revealed no changes to EF across ages or treatments (**Figure 2E**), significant effects were observed in the key diastolic function parameters of EDV, SV, and MV E/A ratio (**Figure 2F-H**). Notably, S100A1 treatment prevents the decline of EDV and SV associated with D2.*mdx* cardiomyopathy, whereas ARC treatment preserves MV E/A ratio. In agreement with these functional benefits of ARC and S100A1 treatment. These data indicate that cardiac expression of both ARC and S100A1 are beneficial for the treatment of D2.*mdx* cardiomyopathy with therapeutically distinct outcomes.

### Bicistronic S100A1-ARC vector improves D2.mdx cardiac phenotype

Due improvements in diastolic function incurred by S100A1 overexpression and the survival benefit of ARC overexpression, we sought to combine these therapeutic transgenes into a single bicistronic vector utilizing an internal ribosomal entry site (IRES)^44^. The canine versions of these transgenes were used in this therapeutic construct, which were also driven by the cTnT promoter and packaged in AAVrh10 vector (**Figure 3A**), to support further translational studies in larger mammals. Systemic treatment of this vector at a dose of 2×10^14^ gc/kg^16, 60^ achieved ∼10-fold elevation of ARC levels and ∼2-fold elevation of S100A1 levels in *mdx* mice that were injected at 1 mo of age (**Figure 3B**). In agreement with the individual transgene experiment, this bicistronic vector significantly reduced IVRT, trended towards a reduction in LV MPI, and significantly increased MV E/A ratio of 12 mo *mdx* mice, while not affecting EF **(Figure 3C**). At 20 mo, the diastolic function measures of LV MPI, MV E/A, MV deceleration time were significantly improved by the S100A1-ARC treatment compared to age-matched controls, whereas EDV and SV both trended towards an increase with treatment without an effect on EF (**Figure 3D**). Interestingly, the lack of significance in S100A1-ARC treated SVs appears to due to increased ESVs in these mice, suggesting that functional reserve of *mdx* hearts may be increased with this treatment. In agreement with these functional improvements, S100A1-ARC treated hearts also exhibit less signs of histopathology than untreated controls (**Figure 3E**). It is important to note that while the S100A1-ARC treatment group underwent longitudinal measures in this study, no mice of the 12 mo control group survived to the 20 mo timepoint. We, therefore, utilized untreated *mdx* mice that survived to this timepoint for age-matched untreated data. These findings indicate that a bicistronic vector consisting of S100A1 and ARC provides significant benefits to the dystrophic myocardium, particularly in regards to improving diastolic function.

**Figure 3.**
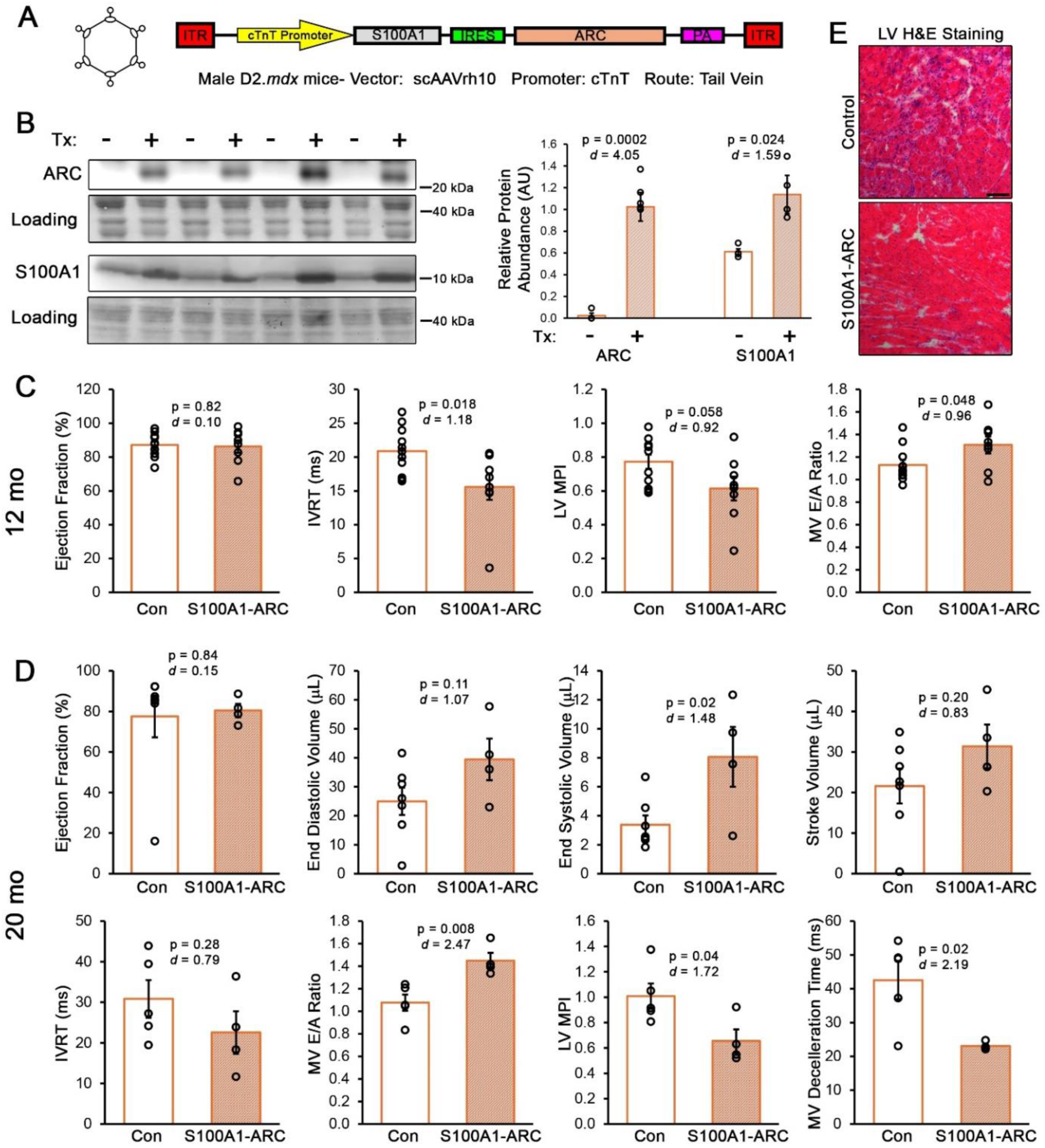
Bicistronic S100A1-ARC provides long-term benefits to D2.*mdx* cardiac function. (**A**) Male D2.*mdx* mice received control (n = 12) or scAAVrh10.cTnT.S100A1-ARC (n = 8) treatments at 1 month of age (mo) and underwent echocardiography at 12 and 20 mo. (**B**) Cardiac ARC and S100A1 protein levels were assessed at study endpoint. Echocardiography measures are shown for both (**C**) 12 mo and (**D**) 20 mo timepoints. (**E**) Representative images of LV histopathology are provided, as analyzed by hematoxylin & eosin (H&E) staining (scale bar represents 100 µm). Data are displayed as mean±SEM with individual values indicated by open circles and were analyzed using two-tailed Welch’s T-test (α = 0.05), with p-values and effect size indicated.

### S100A1-ARC treatment prevents micro-dystrophin associated heart failure

Body-wide micro-dystrophin gene therapies are in the clinic to treat DMD-affected individuals with a miniaturized dystrophin transgene designed to be expressed in all striated muscles^61^. In previous work, we identified that specific micro-dystrophin designs have the ability to promote cardiac dysfunction and premature death in D2.*mdx* mice^16^. This lethality has been confirmed to be caused by HF, as determined by significant increases in the HF markers *Nppb* and *Myh7* and decreases in *Myh6* and the *Myh6/Myh7* ratio relative to untreated *mdx* hearts (**Figure S4**). To evaluate if S100A1-ARC treatment can rescue this phenotype, we treated 1 mo D2.*mdx* mice with a HF-inducing ΔR3-R21 ΔCT micro-dystrophin^16, 62^ alone or in combination with S100A1-ARC (**Figure 4A**). While there was no effect of dual treatment on the expression of micro-dystrophin content, ARC and S100A1 transgene levels were robustly expressed in only the dual vector treatment group (**Figure 4B**). Importantly, S100A1-ARC treatment significantly increased mouse survival from a mean survival time of 336 days to 625 days (**Figure 4C**). Because age-matched micro-dystrophin only mice were not available for comparison of heart function, we compared echocardiography data from 22 mo micro-dystrophin + S100A1-ARC treated mice to those from 20-22 mo WT and 20 mo *mdx* groups. This comparison revealed no significant differences in EF across the groups, whereas SV and EDV from the micro-dystrophin + S100A1-ARC group more closely resembled WT values than those of untreated *mdx* mice (**Figure S5A**). Cardiac histopathology was also markedly improved in the combinational treatment group (**Figure S5B**). These findings demonstrate that the S100A1-ARC bicistronic vector prevents the HF associated with cardiotoxic micro-dystrophin transgenes and may represent an adjunct therapeutic in the event it is found that micro-dystrophin promotes similar forms of HF in the clinic.

**Figure 4.**
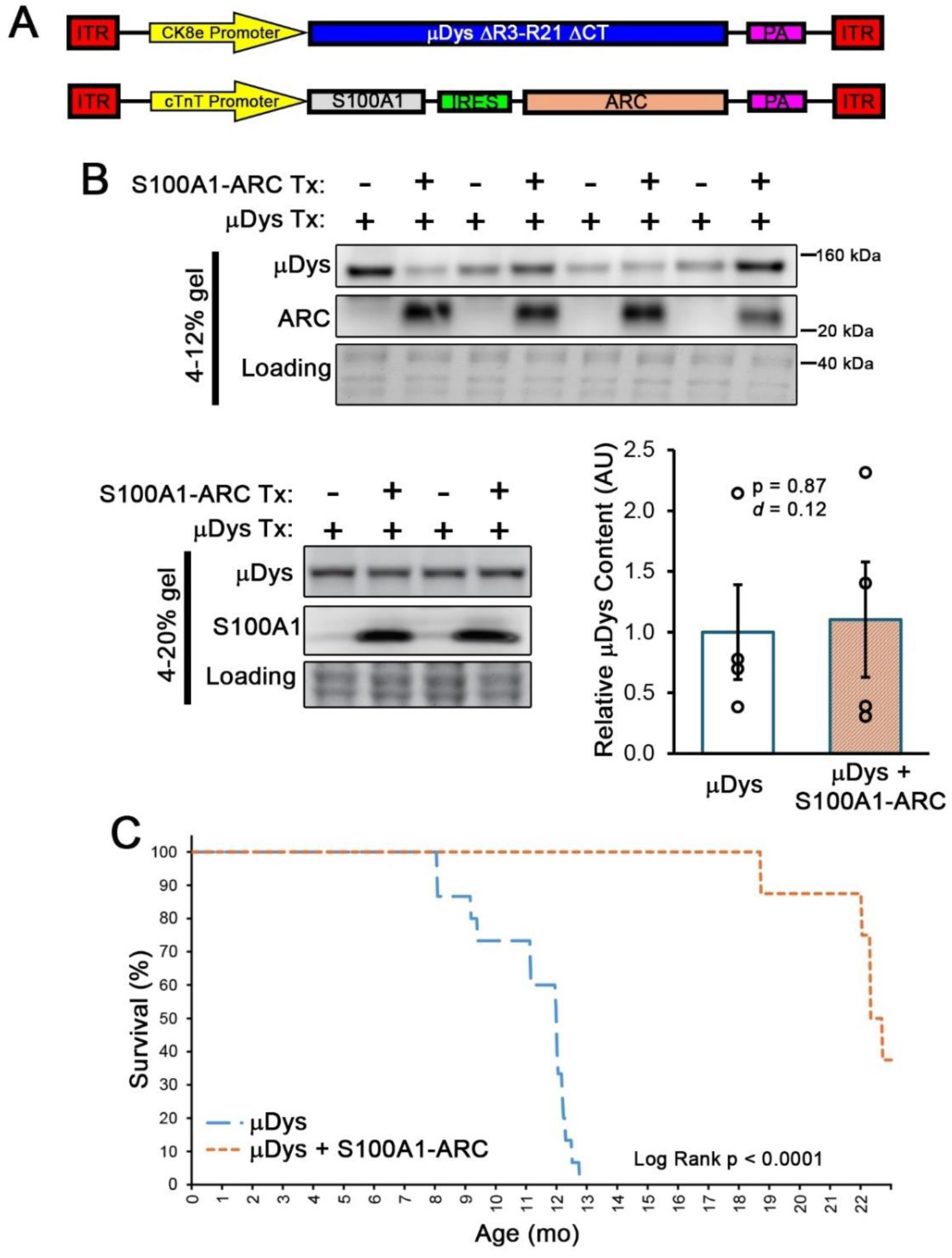
S100A1-ARC gene therapy prevents micro-dystrophin cardiotoxicity. (**A**) Male D2.*mdx* mice were treated with the ΔR3-R21ΔCT micro-dystrophin (µDys) gene therapy with (n = 8) and without (n = 15) co-treatment with the bicistronic cardiac S100A1-ARC vector at 1 month of age (mo) and were evaluated for survival. (**B**) Cardiac expression of µDys, ARC, and S100A1 was measured by immunoblotting at respective humane endpoints of the study. (**C**) Survival analysis reveals significant life-extension by co-treatment with S100A1-ARC. Data are displayed (**B**) as mean±SEM with individual values indicated by open circles and were analyzed using two-tailed Welch’s T-test (α = 0.05), with p-values and effect size indicated, or (**C**) as a Kaplan-Meier survival analysis.

### S100A1-ARC gene therapy safety in GRMD canine model of DMD

We next sought to test the safety of myocardial S100A1-ARC expression in the GRMD canine model of DMD. Prior to S100A1-ARC development, we investigated AAV serotypes capable of efficient transduction of canine hearts via IV administration. While AAV9 exhibited lower transduction than AAV8 in the initial testing (**Figure S6A**), we identified AAVrh10 as a vector that efficiently transduces canine striated muscle at an IV delivered dose of 5×10^12^ gc/kg, compared to AAV8 and AAVrh8 (**Figure S6B-C**). To further reduce amounts of vector required to specifically transduce canine hearts with therapeutic AAVs, we investigated the transduction efficiency of intracoronary AAV delivery, which revealed uniform transduction of the LV, septum, and right ventricle at doses of 2×10^13^ gc (**Figure 5A**) and 5×10^13^ gc (**Figure 5B**). Ensuing therapeutic treatments of S100A1-ARC, therefore, utilize 5×10^13^ gc of vector delivered via intracoronary injections.

**Figure 5.**
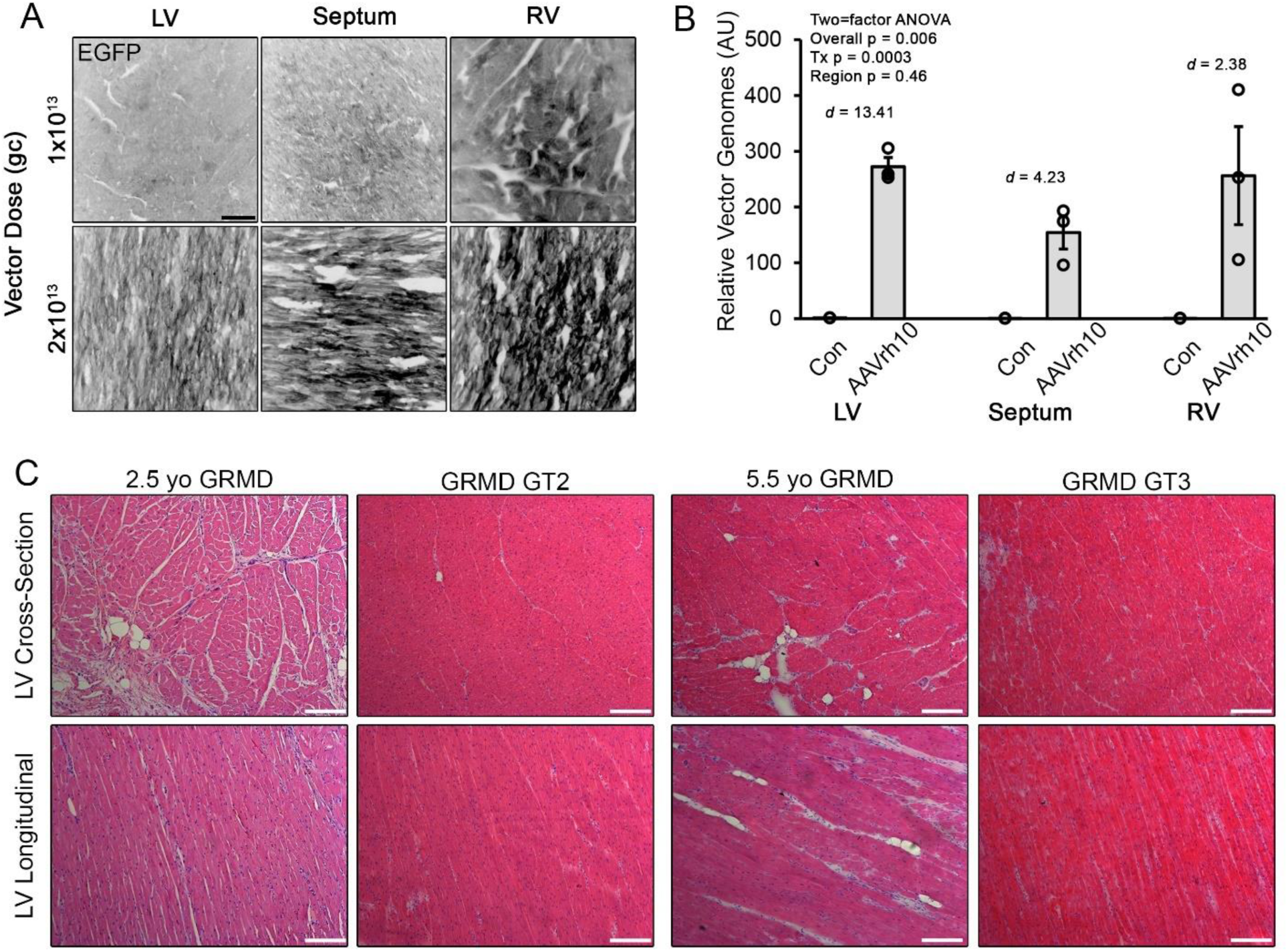
Intracoronary delivery of AAVrh10 in GRMD canines. (**A**) Representative fluorescence images of EGFP expression in the left ventricle (LV), septum, and right ventricle (RV) of canine hearts 1 week following intracoronary delivery of 1×10^13^ or 2×10^13^ gc of AAVrh10.EGFP. (**B**) AAV vector genomes were quantified in the LV, septum, and RV of GRMD hearts following intracoronary treatment with 5×10^13^ gc of AAVrh10 vector. (**C**) Representative hematoxylin & eosin staining of LV cross-section and longitudinal section from S100A1-ARC treated GRMD GT2 and GT3 with comparably age-matched control GRMD samples (scale bar represents 100 µm). Data are displayed as mean±SEM with individual values indicated by open circles and were analyzed using two-factor ANOVA (treatment and region effects). Effect sizes between region-matched groups are indicated.

The bicistronic scAAVrh10.cTnT.S100A1-ARC vector was delivered to three GRMD dogs: GT1 at 5 mo, GT2 at 10 mo, and GT3 at 24 mo. Because GRMD cardiomyopathy is progressive with age^17, 51^, we chose this spectrum of treatment ages to verify safety of vector delivery across ages representing different degrees of preexisting cardiac pathology. All treatments were well tolerated and none of the treated dogs exhibited overt signs of cardiomyopathy during the course of this study, which was followed for years without signs of safety concern from the vector treatment. The longevity and eventual endpoint of these dogs are summarized in **Table S4**.

The low sample size of this safety evaluation limits interpretation of efficacy from this treatment. However, echocardiography data from the treated canines of this group compared to historical age-matched data from the colony revealed large effects sizes for FS improvements at 18-20, 21-24, and 25-30 mo and SV increases at 18-20 mo for GT1 and GT2 (**Figure S7**). In agreement with possible functional benefits associated with these GRMD treatments, tissue sections of GT2 and GT3 LV samples exhibit markedly reduced signs of histopathology compared to comparably age-matched untreated GRMD samples (**Figure 5C**). Together, these data provide evidence of both safety and improved cardiac phenotype by S100A1-ARC gene therapy in the GRMD heart.

### Skeletal muscle function improved by striated muscle expression of S100A1-ARC

Given the robust protection incurred by S100A1-ARC expression in the dystrophic heart, we sought to determine if dystrophic skeletal muscle can also benefit from treatment with this bicistronic transgene, including abrogation of the severe muscle regeneration that occurs with dystrophin loss in muscle fibers. A wave of muscle degeneration occurs early in life for D2.*mdx* mice, beginning at ∼1 mo and peaking at ∼2 mo^40^. Therefore, we investigated systemic treatment of S100A1-ARC driven by the CK8e striated muscle promoter^45^ delivered at 1 mo to coincide with the start of degeneration with an endpoint at 2 mo to gauge functional protection at the peak of skeletal myopathy (**Figure 6A**). At endpoint, the quadriceps of treated mice exhibited robust levels of vector transduction (**Figure 6B**), however skeletal muscle transduction was less than that exhibited in the heart, as previously shown^16, 63^. Importantly, treated EDL muscles exhibit greater maximum force production than controls, whereas the soleus muscles of treated mice demonstrated significantly higher specific tension (tetanic force production normalized to muscle cross-sectional area; **Figure 6C-D**). These data indicate that skeletal muscle functional benefits can be achieved by total striated muscle expression of S100A1-ARC, therefore the benefits of this therapeutic dual transgene are not restricted to the myocardium.

**Figure 6.**
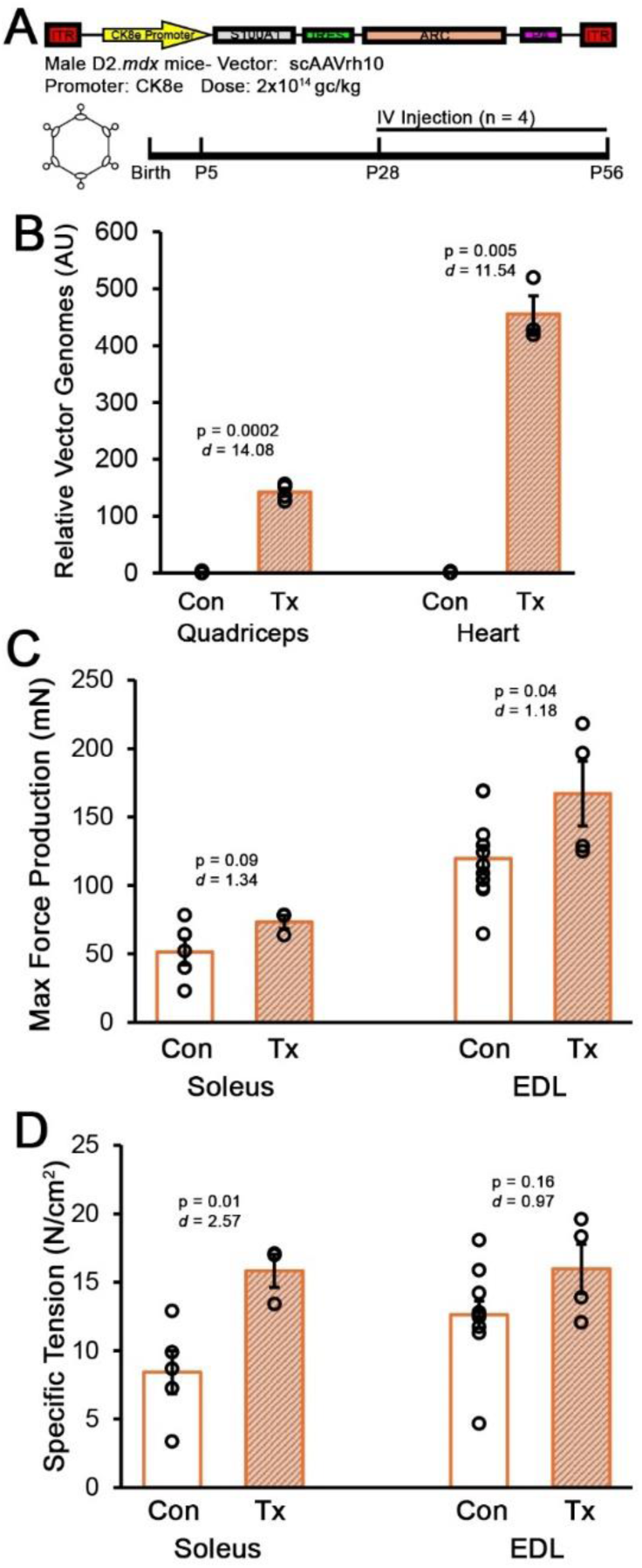
S100A1-ARC provides functional benefits to D2.*mdx* muscle. (**A**) Male D2.*mdx* mice received treatment with control (n = 11) or S100A1-ARC driven by the straited muscle specific CK8e promoter (Tx; n = 4) at 1 month of age (mo) and underwent *ex vivo* functional measures of the soleus and extensor digitorum longus (EDL) muscles at 2 mo. (**B**) Vector genomes were quantified in the quadriceps and hearts from control and treated mice. (**C**) Maximum tetanic force production and (**D**) specific tension from the soleus and EDLs of control and treated D2.*mdx* mice at study endpoint. Data are displayed as mean±SEM with individual values indicated by open circles and were analyzed using two-tailed Welch’s T-test (α = 0.05), with p-values and effect size indicated.

## DISCUSSION

Effective therapeutics for DMD cardiomyopathy are a major unmet clinical need for individuals affected by this devastating disease. Herein, we have identified that AAV-mediated cardiac expression of S100A1 prevents many features of diastolic dysfunction in the dystrophic heart and cardiac ARC overexpression prolongs the life expectancy of D2.*mdx* mice. The combination of the two transgenes in a bicistronic vector improves the functional outcomes and survival of D2.*mdx* hearts, as well as exhibits long-term safety and evidence of cardiac benefits in the GRMD canine model of DMD. Furthermore, S100A1-ARC expression in skeletal muscle reduces functional decrements associated with the early stage of D2.*mdx* muscle disease. Taken together, these findings indicate that the dual expression of S100A1 and ARC in dystrophic hearts represents a promising gene therapy for the treatment of DMD.

In this report, we rigorously demonstrate that progressive diastolic dysfunction and LV restriction are major features of the cardiomyopathy exhibited by the D2.*mdx* murine model of DMD, characterized by pronounced reductions in EDV, SV, and MV E/A. This phenotype, which entail impairments in the LV’s ability to relax and fill with blood before the next contraction, are becoming more recognized as aspects of DMD cardiomyopathy that precede development of systolic dysfunction and DCM^11, 12^. Interestingly, this cardiac phenotype closely resembles HF with preserved EF (HFpEF), a lethal form of cardiomyopathy that commonly affects individuals with metabolic diseases^64^, where reduced EDV and SV results in compensatory increases in heart rate and contractility to maintain adequate CO for survival. Therefore, we suspect this this restrictive phase of DMD cardiomyopathy represents a similar silent form of HF that can occur for a considerable period before progressive myocardial damage and energy deficits lead to sudden death or decompensation to DCM. Therefore, therapeutics effective at treating this phase of DMD cardiomyopathy have the potential to delay onset of DCM, as evidenced by our previous observations using the phosphodiesterase 5 inhibitor, tadalafil^17, 65^.

An important finding in this report is that the mechanism of LV restriction in the dystrophic myocardium appears to be cardiomyocyte-driven, rather than being caused by fibrosis or other extracellular factors. This is evidenced by the acute reversal of reduced EDV by the cardiac myosin inhibitor, mavacamten, and strengthens the hypothesis that DMD cardiomyopathy is largely driven by impaired calcium handling^20–22^. We^16^ and others^66^ have previously identified resting calcium levels to be elevated in dystrophic cardiomyocytes. These elevated calcium levels can lead to a myriad of negative consequences for the cardiomyocyte, including impaired relaxation, E-C coupling abnormalities, mitochondrial dysfunction, aberrant protease activation, and, ultimately, cell death^17, 20, 23, 24, 67–69^. At the organ level, these consequences of cardiomyocyte calcium mishandling result in diastolic dysfunction, life-threatening arrhythmias, and progressive replacement of myocardium with fibrotic scarring, which are all major features of DMD cardiomyopathy^4, 7, 10^.

We sought to address these calcium-associated issues of the dystrophic myocardium by implementing gene therapy to upregulate cardiomyocyte S100A1 and ARC, alone and in combination with each other. While these proteins are both involved in calcium homeostasis^35, 70^, we identified differential benefits incurred by each transgene: S100A1 alleviates LV restriction and ARC prolongs life expectancy. Physiologically, these distinct effects are logical. S100A1 is an EF-hand calcium sensor that can directly buffer calcium, increase SERCA activity, and reduce RyR2 leakage upon activation by calcium binding^35^, thereby reducing cytosolic calcium levels through multiple mechanisms to allow improved myofilament relaxation. ARC can also directly bind calcium^70^, however, it’s primary mechanism of action is to prevent activation of cell death pathways, including apoptosis (intrinsic and extrinsic pathways), necroptosis, and pyroptosis^71^. Because ARC overexpression has previously been shown to decrease cardiomyocyte death in the border zone of post-ischemic myocardium^36^, ARC possibly prevents other forms of cardiomyocyte death, including mechanoptosis^72^. Therefore, ARC ultimately prevents loss of cardiomyocytes and enhances organism survival.

To take advantage of both major benefits incurred by cardiac overexpression of S100A1 and ARC, a bicistronic DNA construct to deliver both transgenes in the same self-complementary AAV vector was developed. Treatment with this dual transgene approach resulted in long-term improvements in D2.*mdx* diastolic function and survivability, HF prevention when combined with a cardiotoxic micro-dystrophin transgene, and long-term safety and histological benefits when delivered to the hearts of GRMD canines. These findings indicate that S100A1-ARC gene therapy represents a promising treatment for DMD cardiomyopathy, which is currently an unmet clinical need for individuals with DMD. From a translational perspective, clinical implementation of this cardiac-directed dual transgene vector is highly feasibility via the intracoronary delivery we used in canines. This delivery requires potentially orders of magnitude less vector than systemic AAV treatments that aim to transducing all musculature^60^, therefore, improves the overall safety profile of AAV delivery, which has been associated with severe and lethal responses in the clinic following systemic delivery with high doses^73, 74^.

Other gene therapy approaches to address DMD cardiomyocyte calcium handling issues primarily involve increasing SERCA activity. This includes directly overexpressing SERCA^26^ and manipulating levels of SERCA modulating proteins^27, 28^. While these approaches do effectively increase SERCA activity, it is unclear how the increased energy requirements of additional active calcium transport impact dystrophic cardiomyocytes, which are already energetically challenged^20^. Furthermore, they do not address the release of calcium from the SR through leaky RyR channels. S100A1 overexpression, on the other hand, effectively addresses these calcium handling issues while also providing direct mitochondrial benefits to increase ATP production in cardiomyocytes^75^. The pleiotropic benefits of S100A1 in cardiomyocyte calcium handling have led to efforts to develop S100A1-based gene therapies for several forms of HF^35^, including a recent report that utilized AAV5-packaged S100A1 to improve functional outcomes in a porcine model of myocardial infarction^76^. To our knowledge, the current work is the first to exploit S100A1’s myriad of benefits in context of DMD cardiomyopathy, which showed promising efficacy at preventing diastolic dysfunction and LV restriction in a severe mouse model of DMD. Particularly in combination with the life-extending benefits of ARC, we anticipate the S100A1-ARC strategy developed in this study to be effective at improving cardiac outcomes in other muscular dystrophies, such as the limb-girdle muscular dystrophies, as well as other forms of cardiomyopathy/HF, including HCM and DCM caused by various genetic and environmental factors.

In addition to the robust cardiac benefits achieved by S100A1-ARC gene therapy, we also provide evidence that this dual-transgene vector provides functional benefits to skeletal muscle when driven by the striated muscle-specific CK8e promoter^45^. As we have previously described^40^, the severe degenerative phase of D2.*mdx* skeletal myopathy initiates at 1 mo and peaks at ∼2 mo. Therefore, the improved muscle function observed in the soleus and EDL following S100A1-ARC treatment was incurred during the most devastating stage of murine muscle disease progression. This was achieved without sarcolemma stabilizing treatments, like micro-dystrophin expression. While additional studies are required to determine optimal dosing and timing of S100A1-ARC as a stand-alone or adjunct gene therapy for skeletal muscle benefits, these data provide proof-of-concept that S100A1-ARC improves muscle function in the early phases of D2.*mdx* skeletal muscle pathology.

The past decade has seen much progress in the field of DMD therapeutics with the regulatory approval of several treatments, including small molecules^77–79^, anti-sense oligonucleotides^80^, and AAV gene therapy^61^. Despite these advancements in therapeutics focused on skeletal muscle outcomes, DMD cardiomyopathy is still primarily managed by treatment with beta-blockers, ACE inhibitors/angiotensin receptor blockers, and/or mineralcorticoid receptor agonists^81, 82^. In this report, we identify S100A1-ARC gene therapy as an effective treatment for DMD cardiomyopathy through rigorous efficacy testing in D2.*mdx* mice and a safety evaluation in the GRMD large animal model of DMD. This discovery is very timely, as systemic micro-dystrophin gene therapy is now approved for DMD, and it is currently unknown how these miniaturized dystrophin transgenes modify the progression of DMD cardiomyopathy in the clinic. The fact that high cardiac expression of some micro-dystrophin transgenes can induce premature HF and death in mice is very concerning^16^; however, it is also unclear what expression levels are being experienced in the human heart following gene therapy. Nonetheless, our findings indicate that S100A1-ARC therapy represents a novel and promising treatment strategy for DMD cardiomyopathy that can be administered alone or in combination with micro-dystrophin and, likely, other emerging dystrophin-replacement strategies. Furthermore, the dystrophin-independent nature of this therapeutic approach conveys a high probability of also being advantageous for the treatment of other muscular dystrophies, myopathies, and cardiomyopathies.

## ACKNOWLEDGEMENTS AND FUNDING SOURCES

The authors greatly thank Eli Zerpa, Lauren Weigle, Brooke Seldin, Anton Ramick, Victor Prima, and Orinda Hobson for their technical assistance in this work, as well as Michael Matheny, Aimee Goulet, and members of the University of Florida Physiological Assessment Core for facilitating animal work associated with this project. Funding for this project was provided by an NIH/NIAMS Wellstone Center grant (P50-AR052646) to HLS and DWH, a Parent Project Muscular Dystrophy grant to HLS, a Leducq Foundation grant (13CVD04) to HLS, a Department of Defense DMDRP Idea Development grant (W81XWH-22-1-1070) to DWH, and an NIH/NIAMS grant (R01-AR083108) to DWH.

## SUPPLEMENTAL MATERIALS

## SUPPLEMENTAL METHODS

### Intracoronary vector delivery

Metoclopromide (0.5 mg/kg) and maropitant citrate (1 mg/kg) were administered subcutaneously prior to the procedure and an intravenous catheter was placed routinely. Dogs were induced with propofol (2-10 mg/kg) intravenously. Dogs were intubated and anesthesia was maintained with isoflurane (1-3%) with additional propofol and fentanyl (0.7 mcg/kg/min) as necessary. Following induction dogs were placed in right lateral recumbency and the right groin region was clipped of fur and aseptically prepared in a routine manner. A skin incision was performed over the right femoral artery using a scalpel blade. A 6 Fr introducer was placed using the modified Seldinger technique. A 5 French pigtail catheter was inserted through the introducer and advanced into the aortic root. A contrast study was performed with 1 mL/kg of iodinated contrast using a power injector to highlight coronary arteries. Various coronary catheters were selected based on the coronary anatomy observed on the angiogram (4 French diagnostic coronary catheters). Small volumes of diluted contrast were injected during the procedure to ensure good seating of the coronary catheter in the coronary ostium. A total dose of 1-5×10^13^ genome copies of vector (up to 5 mL volume) was administered with 2/3 of the volume administered into the left coronary and 1/3 of the total volume administered into the right coronary. Fifteen seconds before vector administration into each coronary artery, a CRI of adenosine was initiated at a dose of 1 mg/kg/min. The vector was then injected over 15 seconds and the adenosine infusion was continued for an additional 30 seconds. After administration in both coronary arteries, the catheter was removed. The femoral artery was ligated proximal to the catheter insertion site with 2-0 silk two circumferential sutures, and one circumferential suture placed distally. The incision was closed in three layers (muscle, subcutaneous, and intradermal) in a simple continuous pattern using 3-0 PDS.

### Mouse echocardiography and electrocardiograms

Mice were anesthetized using 3% isoflurane and maintained at 1.5-2% to keep heart and respiration rates consistent among treatment groups. Body temperature was maintained at 37℃ throughout imaging. Electrocardiograms were imported into LabCharts (ADInstruments) for analysis. Four images were acquired for each animal: B-mode parasternal long axis (LAX), B-mode short axis (SAX), M-mode SAX, and apical four-chamber view with color doppler and pulsed-wave doppler. M-mode SAX images were acquired at the level of the papillary muscle. Flow through the mitral valve was sampled at the point of highest velocity, as indicated by aliasing, with the pulsed-wave angle matching the direction of flow. Images were imported into Vevo LAB for analysis. Measurements of M-mode SAX and pulsed-wave doppler images were made from three consecutive cardiac cycles between respirations.

### Mavacamten treatment

For the acute mavacamten treatment experiment, mavacampten/MYK-461 (Selleckchem# S8861) was diluted in cherry syrup vehicle (Humco# 00395266216) and administered to mice per oral (PO) at a dose of 2.5 mg/kg.

## SUPPLEMENTAL TABLES

**Table S1.** Electrocardiogram measures from D2.WT and D2.*mdx* mice.

| Age (mo) | Genotype | RR Interval (ms) | Heart Rate (BPM) | PR Interval (ms) | P Duration (ms) |
| --- | --- | --- | --- | --- | --- |
| 2 | WT | 146.68±5.64 | 411.36±14.49 | 39.76±1.26 | 7.95±0.35 |
|  | *mdx* | 143.10±5.01 | 422.98±13.22 | 37.61±1.63 | 8.04±0.21 |
| 3 | WT | 132.73±5.44 | 455.45±16.55 | 34.46±0.86 | 8.19±0.97 |
|  | *mdx* | 139.15±4.09 | 433.10±13.18 | 33.96±0.78 | 7.60±0.33 |
| 4 | WT | 134.84±2.80 | 445.76±9.52 | 34.76±1.36 | 6.94±0.22 |
|  | *mdx* | 136.85±2.14 | 439.44±6.50 | 35.07±0.95 | 7.87±0.27 |
| 6 | WT | 139.19±4.01 | 435.36±11.58 | 36.12±0.95 | 7.51±0.37 |
|  | *mdx* | 151.59±5.58 | 400.44±14.06 | 35.84±0.85 | 8.01±0.71 |
| 8 | WT | 135.09±3.77 | 446.34±11.28 | 34.89±0.99 | 7.75±0.40 |
|  | *mdx* | 141.03±2.23 | 426.25±6.68 | 34.43±1.18 | 8.24±0.72 |
| 10 | WT | 138.78±3.94 | 435.30±11.61 | 34.36±1.12 | 7.84±0.34 |
|  | *mdx* | 141.00±1.85 | 426.01±5.71 | 35.17±1.10 | 7.79±0.46 |
| 12 | WT | 139.30±3.99 | 432.77±11.83 | 33.10±1.13 | 8.06±0.36 |
|  | *mdx* | 140.84±3.31 | 427.63±10.08 | 34.84±0.62 | 7.59±0.63 |
| ANOVA<br>P-values | Overall | 0.21 | 0.13 | 0.06 | 0.98 |
|  | Genotype | 0.59 | 0.53 | 0.23 | 0.91 |
|  | Age | 0.12 | 0.14 | **0.0033** | 0.94 |

Data are presented as mean±SEM and were analyzed using two-factor ANOVA (genotype and age effects) followed by Tukey post-hoc tests ( $\alpha = 0.05$ ). \* $p < 0.05$ vs. age-matched WT values.

**Table S1 Continued**
| Age (mo) | Genotype | QRS Interval (ms) | QT Interval (ms) | QTc Interval (ms) | JT Interval (ms) |
| --- | --- | --- | --- | --- | --- |
| 2 | WT | 13.47±1.53 | 17.56±1.61 | 45.81±3.94 | 4.09±0.85 |
|  | *mdx* | 11.29±0.37 | 20.89±0.77 | 55.36±2.06 | 9.61±0.81* |
| 3 | WT | 9.84±0.41 | 16.18±0.62 | 44.52±1.82 | 6.34±0.84 |
|  | *mdx* | 10.32±0.30 | 17.84±0.35 | 47.93±1.38 | 7.52±0.23 |
| 4 | WT | 10.47±0.89 | 15.91±0.53 | 43.37±1.56 | 5.44±0.48 |
|  | *mdx* | 10.74±0.22 | 19.17±0.52 | 51.83±1.28 | 8.43±0.49 |
| 6 | WT | 10.37±0.37 | 16.63±0.47 | 44.77±1.44 | 6.27±0.60 |
|  | *mdx* | 10.30±0.50 | 19.51±0.46* | 50.18±0.74 | 9.21±0.51* |
| 8 | WT | 10.48±0.74 | 16.12±0.77 | 44.07±2.40 | 5.64±0.57 |
|  | *mdx* | 10.13±0.47 | 19.12±0.44 | 50.99±1.39 | 8.99±0.47* |
| 10 | WT | 10.25±0.71 | 18.00±0.66 | 48.46±1.96 | 7.75±0.54 |
|  | *mdx* | 10.43±0.67 | 19.78±0.44 | 52.66±1.04 | 9.34±0.54 |
| 12 | WT | 9.85±0.92 | 16.95±0.47 | 45.43±1.01 | 7.09±0.98 |
|  | *mdx* | 10.24±0.44 | 20.82±0.68* | 55.61±2.12 | 10.59±0.65* |
| ANOVA P-values | Overall | 0.10 | **<0.0001** | **<0.0001** | **<0.0001** |
|  | Genotype | **0.023** | **0.0014** | **0.0010** | **<0.0001** |
|  | Age | **0.0090** | **0.013** | 0.080 | **0.013** |
|  | Genotype*Age | 0.44 | 0.63 | 0.44 | 0.077 |

Data are presented as mean±SEM and were analyzed using two-factor ANOVA (genotype and age effects) followed by Tukey post-hoc tests ( $\alpha = 0.05$ ). \*p < 0.05 vs. age-matched WT values.

**Table S1 Continued**
| Age (mo) | Genotype | T<sub>peak</sub> T<sub>end</sub> Interval (ms) | P Amplitude (V) | Q Amplitude (V) |
| --- | --- | --- | --- | --- |
| 2 | WT | 2.63±0.62 | 108.04±14.35 | 4.19±3.11 |
|  | *mdx* | 6.68±0.69* | 176.46±14.81* | 18.96±3.05 |
| 3 | WT | 3.83±0.49 | 127.32±10.47 | 13.46±3.72 |
|  | *mdx* | 4.76±0.22 | 148.53±15.68 | 11.64±2.72 |
| 4 | WT | 3.50±0.36 | 112.89±11.76 | 14.45±3.76 |
|  | *mdx* | 5.74±0.44 | 149.40±6.52 | 16.51±3.53 |
| 6 | WT | 3.96±0.40 | 117.02±10.37 | 13.23±3.50 |
|  | *mdx* | 6.31±0.76 | 104.16±8.59 | 16.31±2.47 |
| 8 | WT | 3.97±0.58 | 103.24±9.71 | 6.53±3.60 |
|  | *mdx* | 6.49±0.59 | 98.97±14.72 | 17.38±7.30 |
| 10 | WT | 5.60±0.59 | 104.22±7.37 | 16.08±5.02 |
|  | *mdx* | 6.15±0.64 | 98.56±9.42 | 12.66±12.15 |
| 12 | WT | 4.72±0.65 | 112.73±4.97 | 15.42±2.84 |
|  | *mdx* | 7.06±0.63 | 103.85±13.64 | 5.30±8.18 |
| ANOVA<br>P-values | Overall | **<0.0001** | **<0.0001** | 0.75 |
|  | Genotype | **<0.0001** | **0.0002** | 0.078 |
|  | Age | 0.084 | **0.0006** | 0.96 |
|  | Genotype*Age | 0.15 | **0.0056** | 0.33 |

Data are presented as mean±SEM and were analyzed using two-factor ANOVA (genotype and age effects) followed by Tukey post-hoc tests ( $\alpha = 0.05$ ). \*p < 0.05 vs. age-matched WT values.

**Table S1 Continued**
| Age (mo) | Genotype | R Amplitude (V) | S Amplitude (V) | T Amplitude (V) |
| --- | --- | --- | --- | --- |
| 2 | WT | 526.58±82.98 | -271.39±66.80 | 97.33±24.77 |
|  | *mdx* | 640.01±31.32 | -434.63±62.74 | 316.12±36.68* |
| 3 | WT | 574.45±49.26 | -276.58±81.13 | 217.69±53.93 |
|  | *mdx* | 516.27±41.67 | -381.04±84.66 | 298.87±23.57 |
| 4 | WT | 598.22±103.36 | -262.56±135.35 | 235.32±40.83 |
|  | *mdx* | 648.39±60.52 | -478.20±89.14 | 316.32±33.01 |
| 6 | WT | 510.52±39.85 | -245.54±53.24 | 214.28±26.84 |
|  | *mdx* | 566.08±50.45 | -412.47±121.80 | 318.25±35.20 |
| 8 | WT | 526.98±50.96 | -143.59±64.31 | 218.69±38.14 |
|  | *mdx* | 495.33±49.27 | -96.30±73.77 | 291.89±33.75 |
| 10 | WT | 487.88±63.63 | -115.00±59.98 | 276.64±27.12 |
|  | *mdx* | 506.91±52.09 | -386.66±126.10 | 401.71±48.10 |
| 12 | WT | 496.76±42.21 | -61.00±47.21 | 228.85±46.46 |
|  | *mdx* | 554.03±36.20 | -551.79±73.68* | 398.49±36.33 |
| ANOVA P-values | Overall | 0.4 | **0.0002** | **<0.0001** |
|  | Genotype | 0.19 | 0.23 | **0.0003** |
|  | Age | 0.33 | 0.077 | **0.028** |
|  | Genotype*Age | 0.82 | 0.11 | 0.47 |

Data are presented as mean±SEM and were analyzed using two-factor ANOVA (genotype and age effects) followed by Tukey post-hoc tests ( $\alpha = 0.05$ ). \*p < 0.05 vs. age-matched WT values.

**Table S2.** Heart and cardiomyocyte size in 12 mo D2.WT and D2.*mdx* mice.

| Genotype | WT | *mdx* | P-value |
| --- | --- | --- | --- |
| Heart Mass (mg) | 196.96±3.96 | 146.24±5.38 | 0.0000000052 |
| Heart Mass:TL (mg/mm) | 10.81±0.40 | 8.26±0.28 | 0.0000053 |
| LV Length (mm) | 8.34±0.07 | 7.90±0.16 | 0.019 |
| Cardiomyocyte MFD (µm) | 24.36±1.19 | 17.23±1.86 | 0.022 |

Data are presented as mean±SEM and were analyzed using two-tailed Welch's T-test ( $\alpha = 0.05$ )
TL- tibia length
LV- Left ventricle
MFD- minimum Feret diameter

**Table S3.**
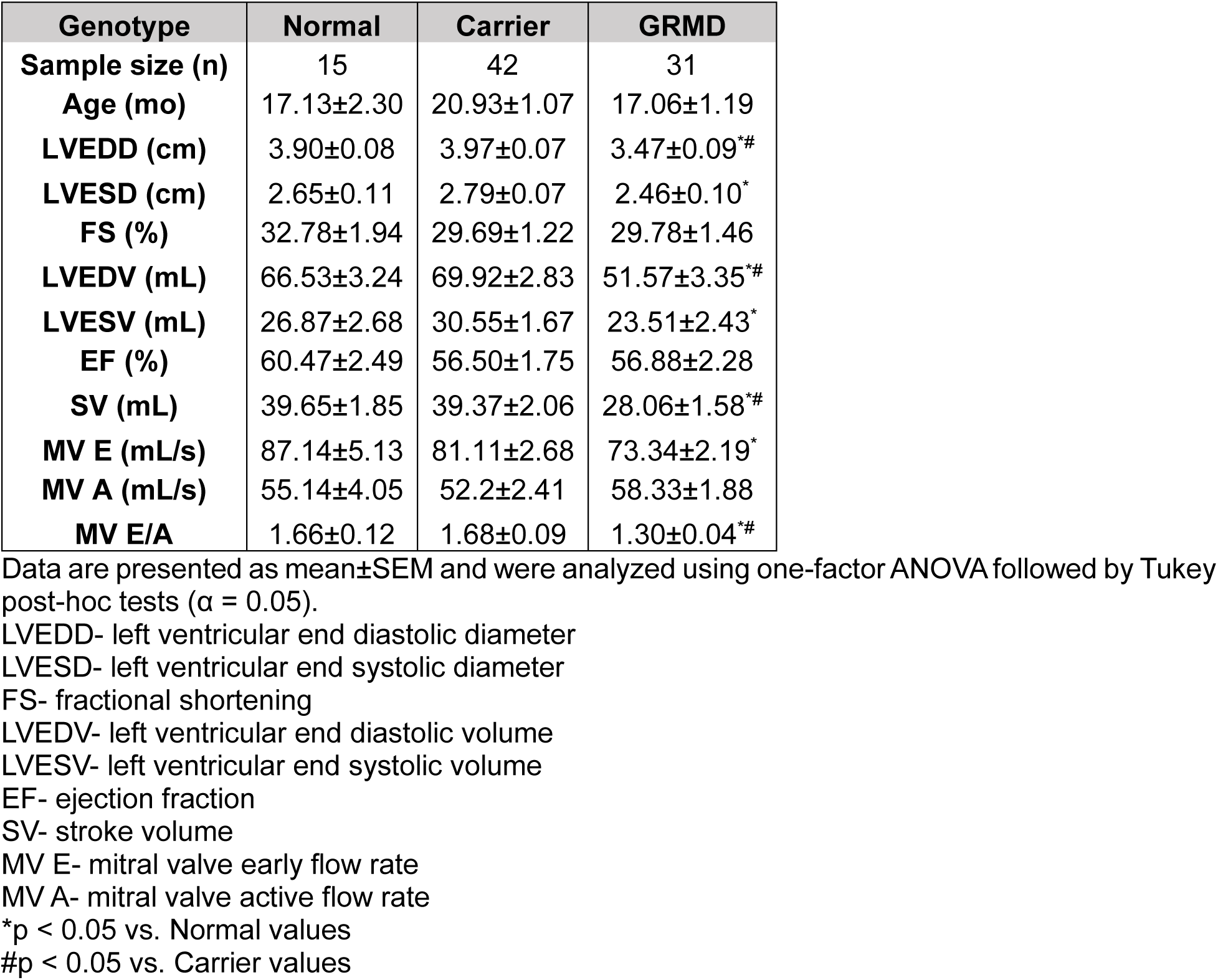
LV Restriction in Pre-DCM GRMD canines.

**Table S4.** GRMD gene therapy summary.

| Subject | Age of Tx (mo) | Age of Death (yrs) | Endpoint Criteria |
| --- | --- | --- | --- |
| GRMD GT1 | 5 | 3.9 | GI torsion |
| GRMD GT2 | 10 | 2.9 | Pneumonia |
| GRMD GT3 | 24 | 5.5 | End of study |

## SUPPLEMENTAL FIGURES

**Figure S1.**
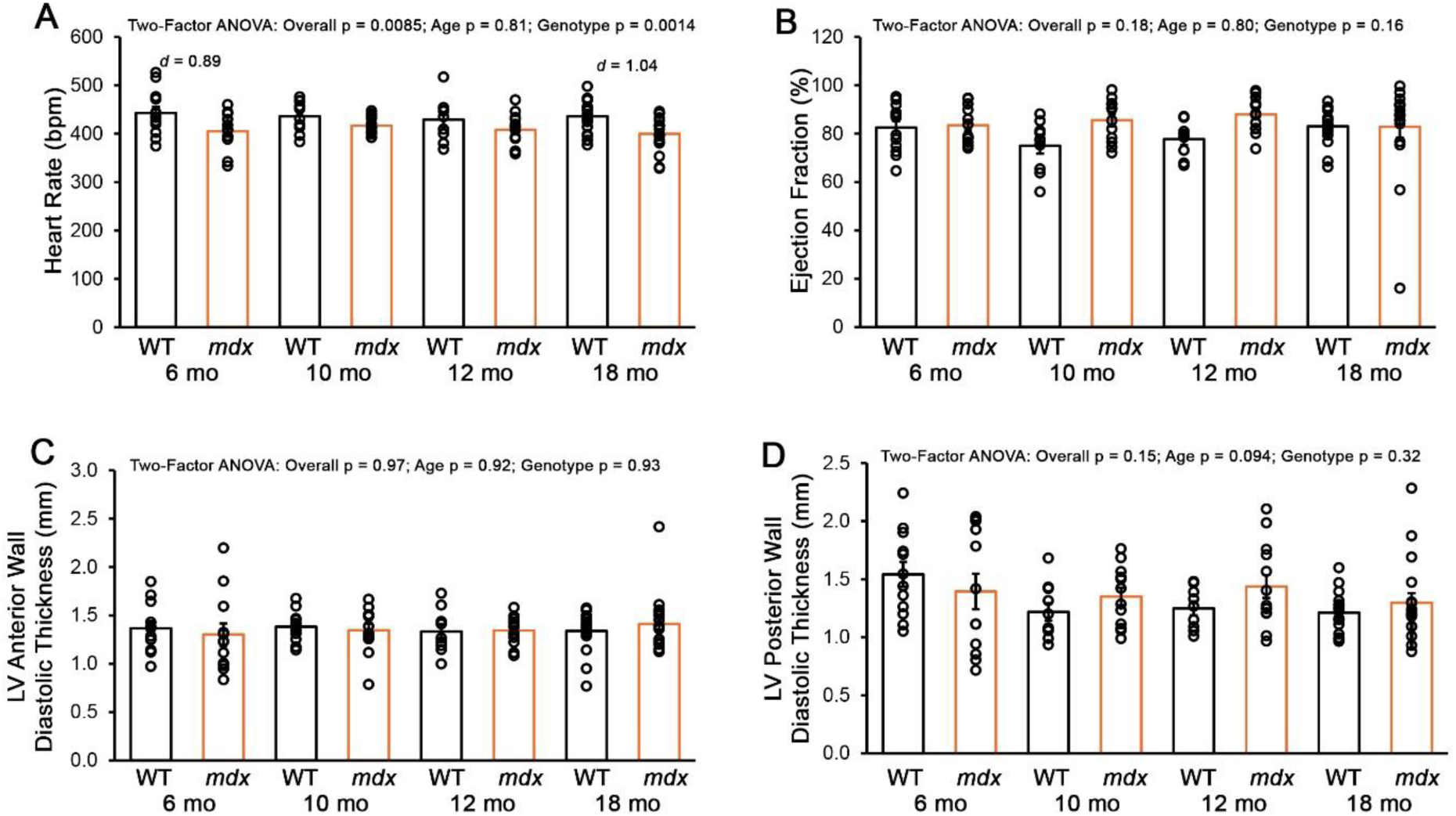
Additional echocardiography measures of D2.WT and D2.*mdx* mice. (**A**) Heart rate, (**B**) ejection fraction, (**C**) diastolic LV anterior wall thickness, and (**D**) diastolic LV posterior wall thickness in 6, 10, 12, and 18 month-old D2.WT (WT) and D2.*mdx* (*mdx*) mice. Data are displayed as mean±SEM with individual values indicated by open circles and were analyzed using two-factor ANOVA (age and genotype effects) followed by Tukey post-hoc tests (α = 0.05). Large effect sizes between age-matched genotypes are also indicated.

**Figure S2.**
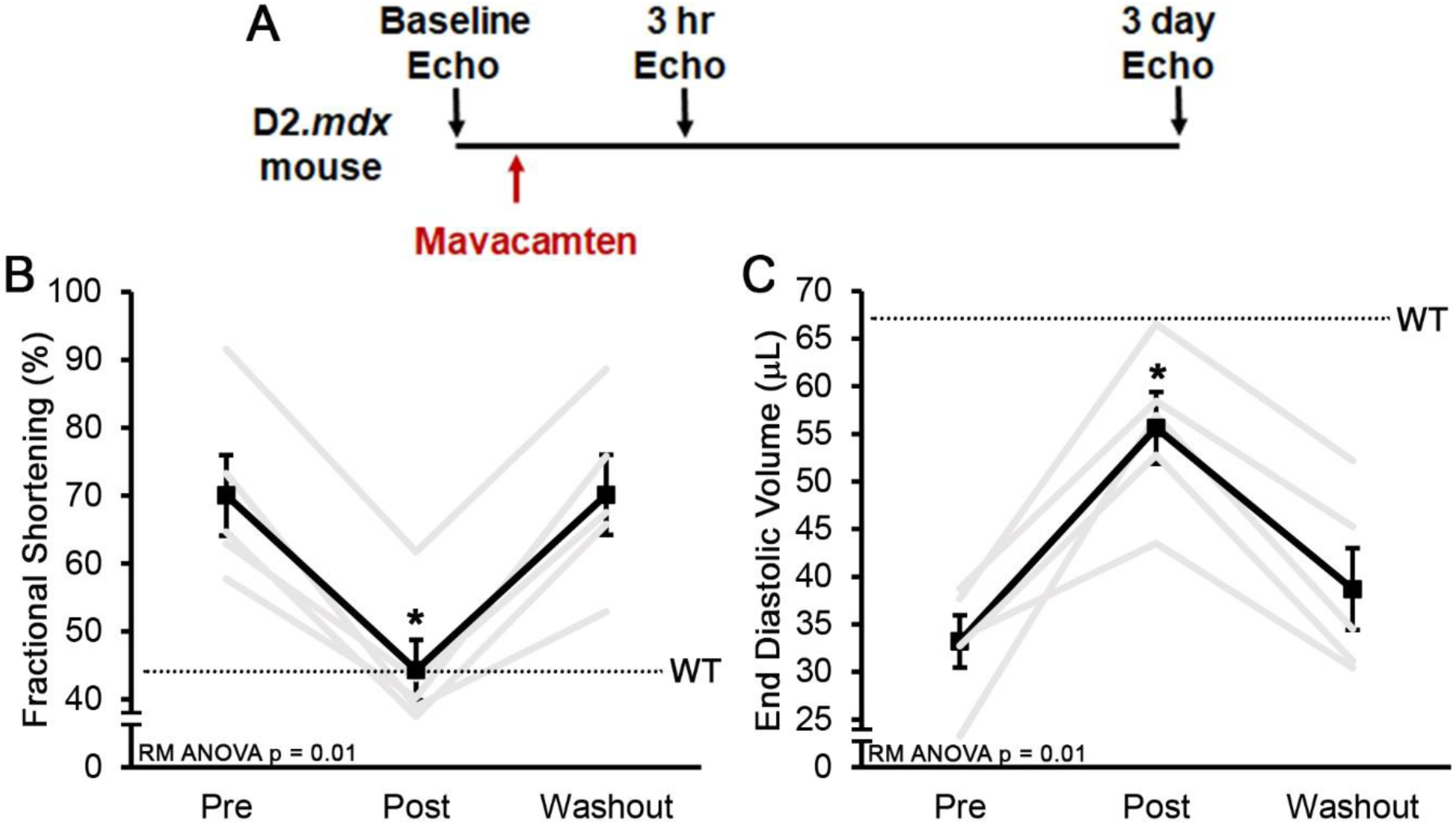
Mavacamten acutely reverses left ventricular restriction in D2.*mdx* mice. (**A**) Ten month-old male D2.*mdx* mice (n = 5) underwent baseline echocardiography (echo) to confirm onset of restrictive phenotype, as determined by low end diastolic volume. Following recovery from anesthesia, mice received an oral dose of 2.5 mg/kg mavacamten. Mice underwent a second echo 3 hours following mavacamten dosing, then a third echo after a 3 day washout period. (**B**) Fractional shortening and (**C**) end diastolic volume from this experiment are depicted as mean±SEM (black line) with individual trajectories (gray lines). Data were analyzed using repeated measures (RM) ANOVA followed by Tukey post-hoc tests (α = 0.05; *p <0.05 vs. pre-treatment values).

**Figure S3.**
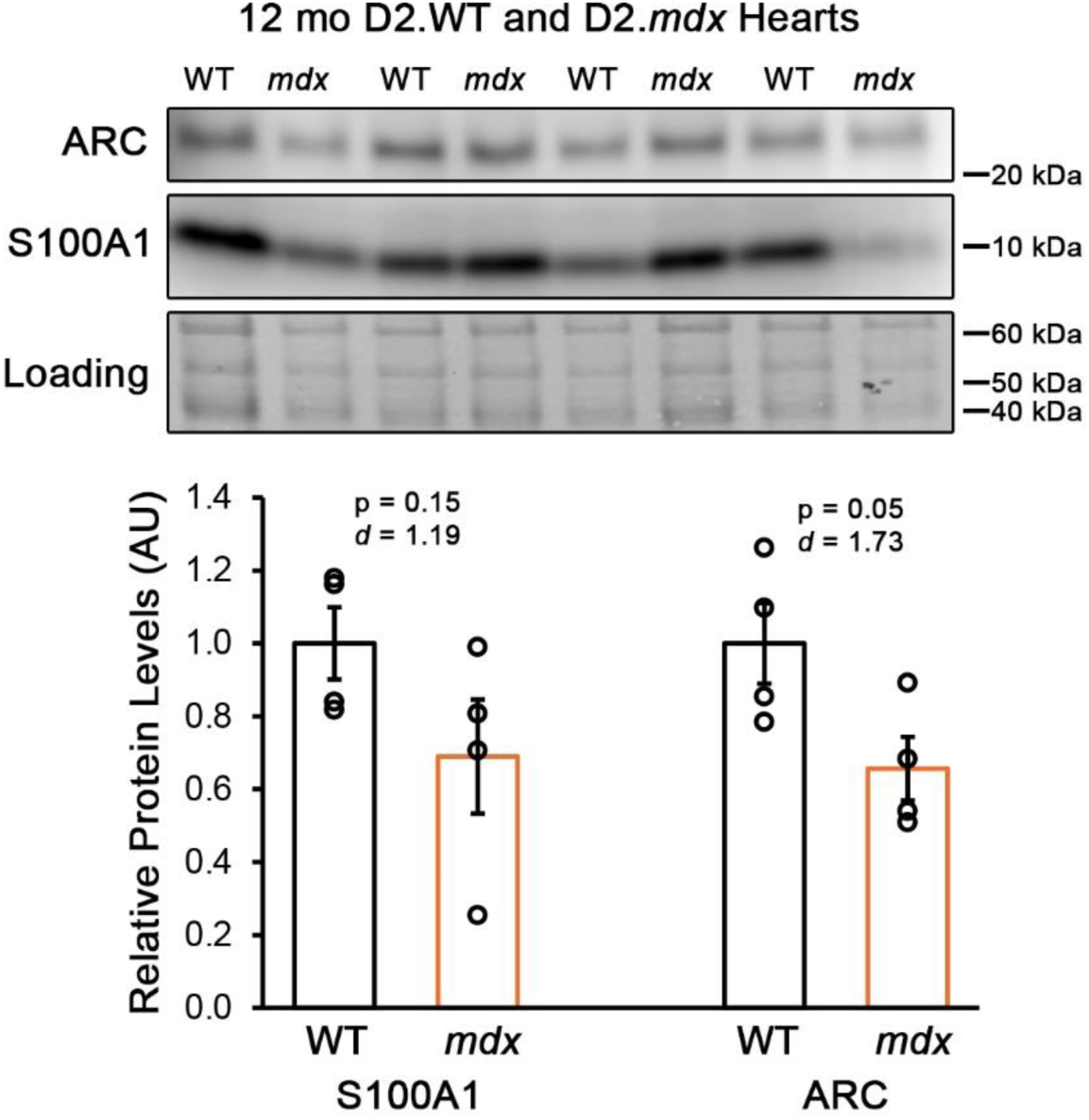
S100A1 and ARC levels in D2.WT and D2.*mdx* hearts. Immunoblotting data for S100A1 and ARC in 12 month-old (mo) D2.WT (WT) and D2.*mdx* (*mdx*) hearts. Data are displayed as mean±SEM with individual values indicated by open circles and were analyzed using two-tailed Welch’s T-test (α = 0.05) with effect size indicated.

**Figure S4.**
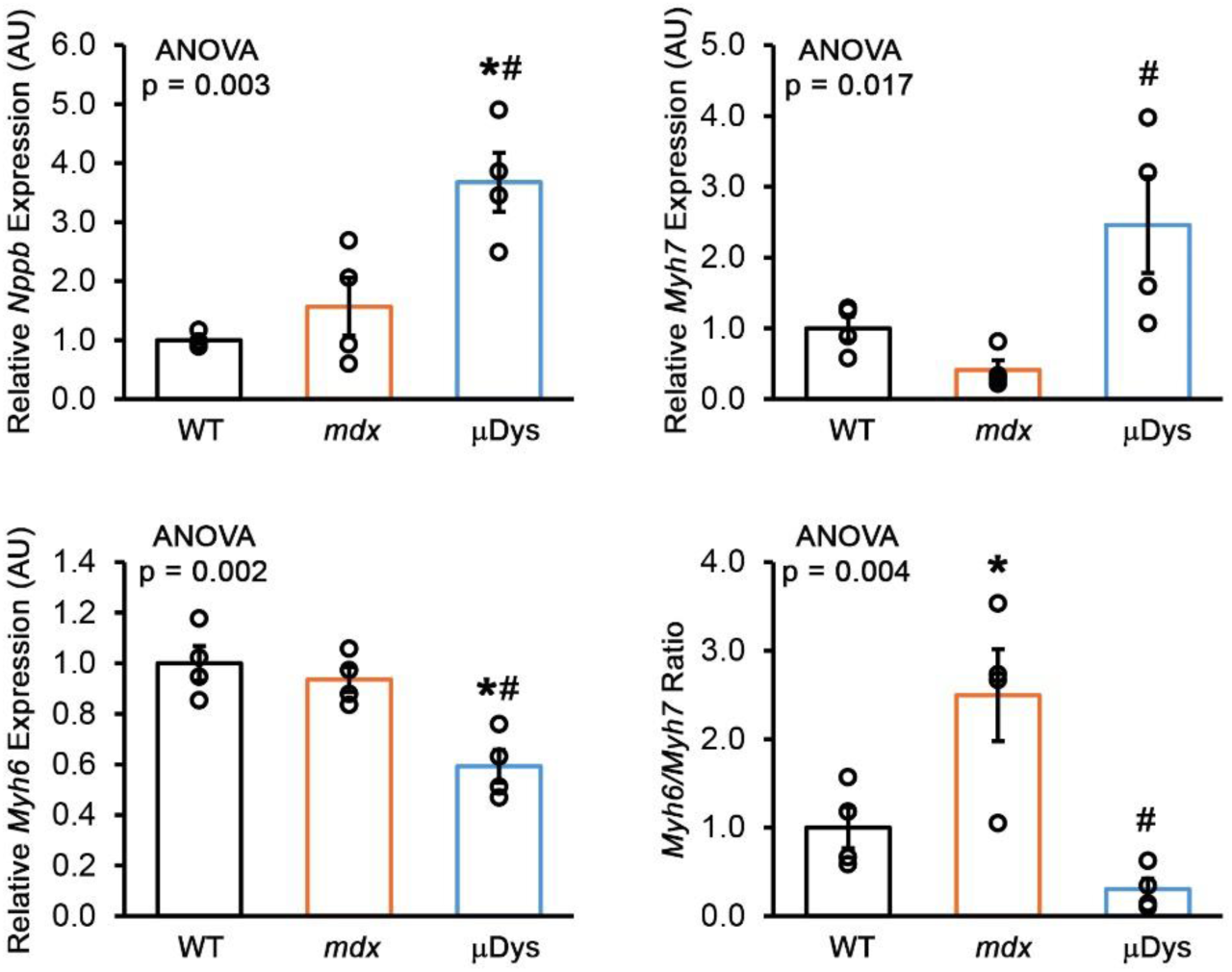
Heart failure associated markers in micro-dystrophin treatment D2.*mdx* hearts. Gene expression of *Nppb*, *Myh7*, and *Myh6*, as well as the *Myh6/Myh7* ratio in 12 month-old D2.WT (WT; n = 4), D2.*mdx* (*mdx*; n = 4), and D2.*mdx* mice that received systemic treatment with the ΔR3-R21ΔCT micro-dystrophin (µDys; n = 4; all reached humane endpoints). Data are displayed as mean±SEM with individual values indicated by open circles and were analyzed using one-factor ANOVA followed by Tukey post-hoc tests (α = 0.05; *p <0.05 vs. WT values; #p <0.05 vs. *mdx* values).

**Figure S5.**
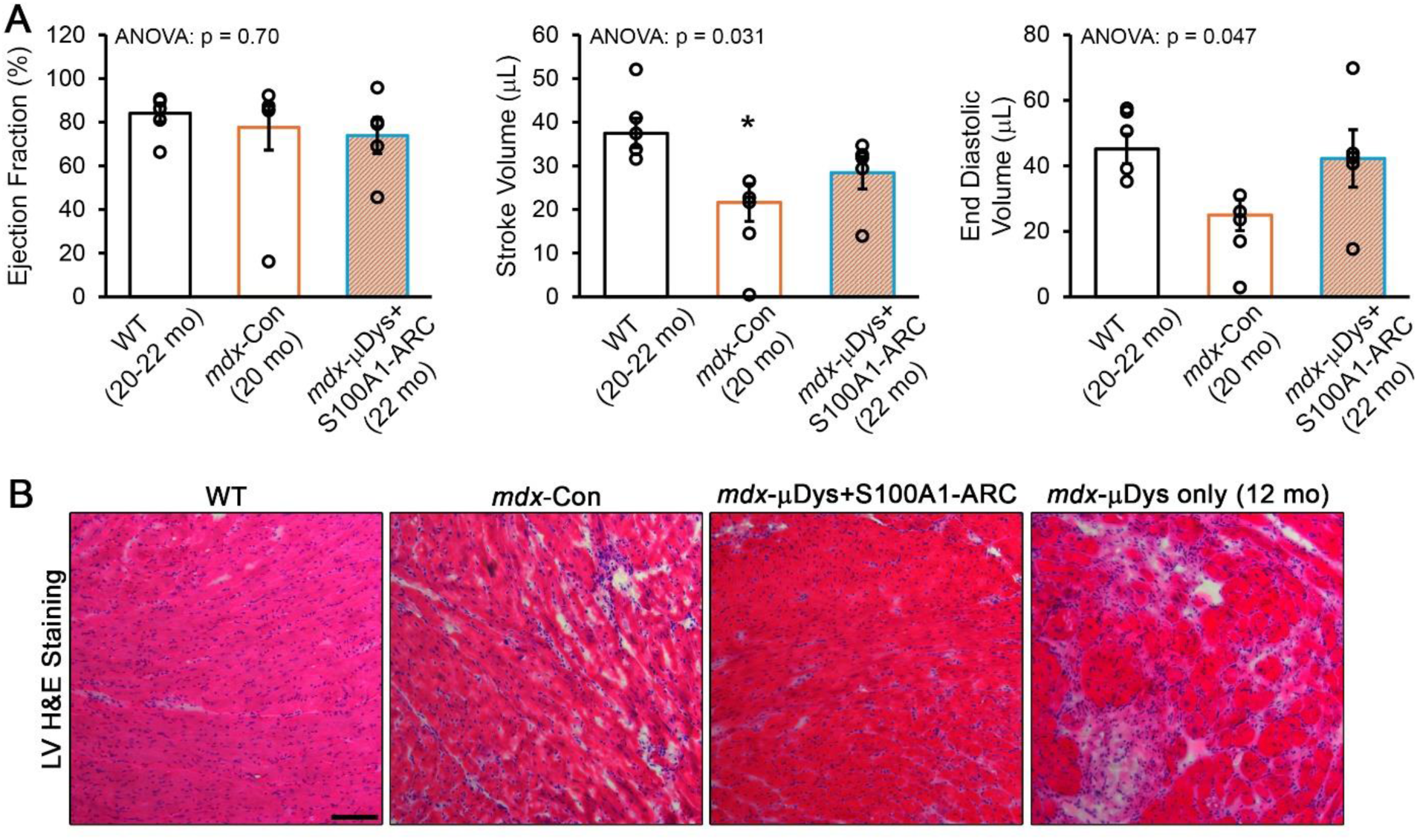
Cardiac phenotypes in mice treated with S100A1-ARC and micro-dystrophin. (**A**) Echocardiographic measures of ejection fraction, stroke volume, and end diastolic volume in 20-22 month-old (mo) D2.WT (WT; n = 6), 20 mo control D2.*mdx* (*mdx-*Con; n = 7), and 22 mo D2.*mdx* that received both micro-dystrophin and S100A1-ARC treatments at 1 mo (*mdx*-µDys+S100A1-ARC; n = 5). (**B**) Representative images of left ventricular (LV) histopathology in WT, *mdx*-Con, *mdx*-µDys+S100A1-ARC, and 12 mo *mdx*-µDys only (reached humane endpoint), as assessed with hematoxylin & eosin (H&E) staining (scale bar represents 100 µm). Data are displayed as mean±SEM with individual values indicated by open circles and were analyzed using one-factor ANOVA followed by Tukey post-hoc tests (α = 0.05; *p <0.05 vs. WT values).

**Figure S6.**
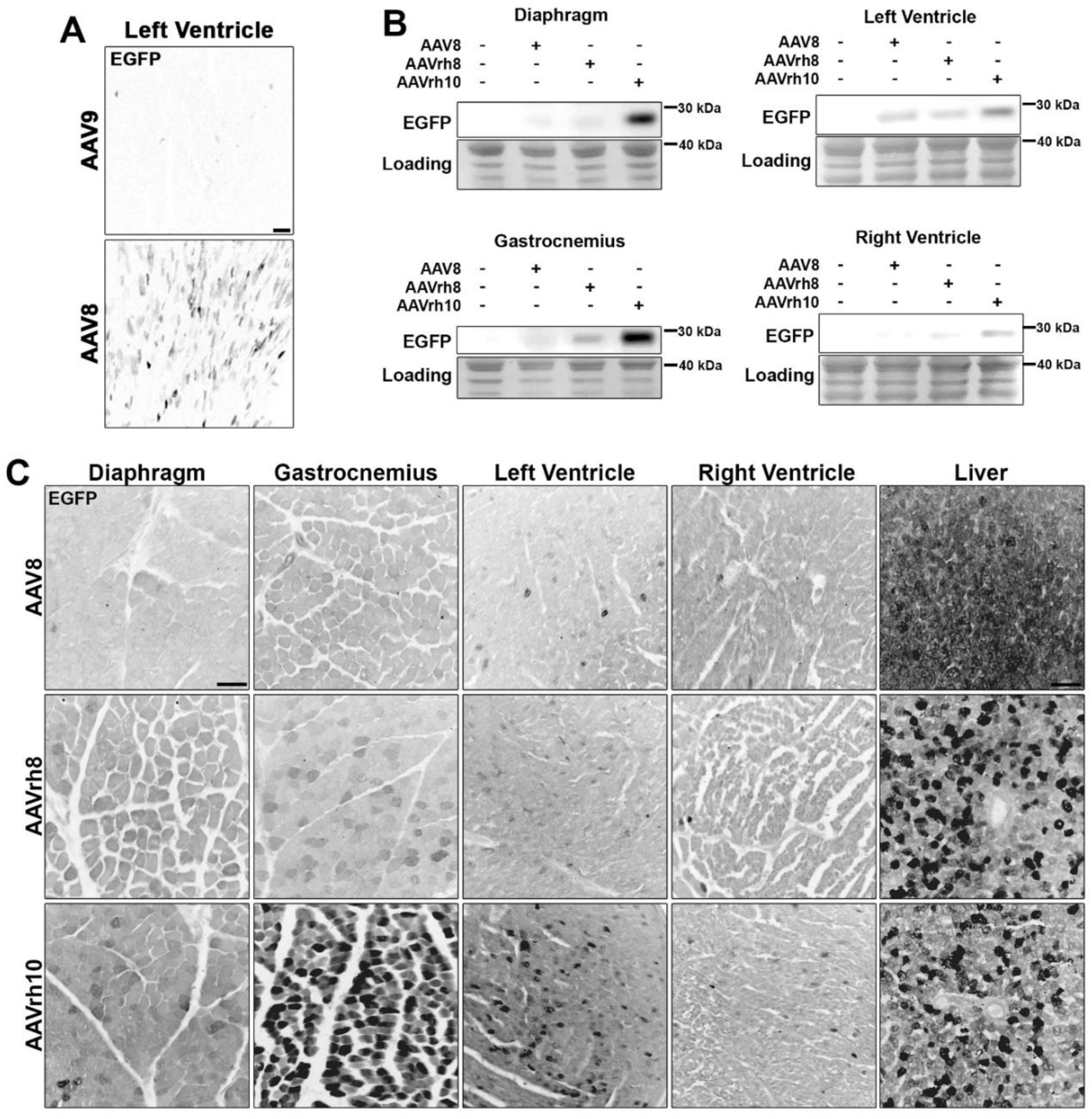
Biodistribution of gene therapy vectors in canine striated muscle. (**A**) Representative EGFP fluorescence in the left ventricles of canines 1 week following intravenous administration of AAV8 or AAV9 at a dose of 5×10^12^ gc/kg. (**B**) Immunoblotting and (**C**) representative EGFP fluorescence in canine tissue 1 week following intravenous administration of AAV8, AAVrh8, or AAVrh10 at a dose of 5×10^12^ gc/kg. Scale bars represent 100 µm.

**Figure S7.**
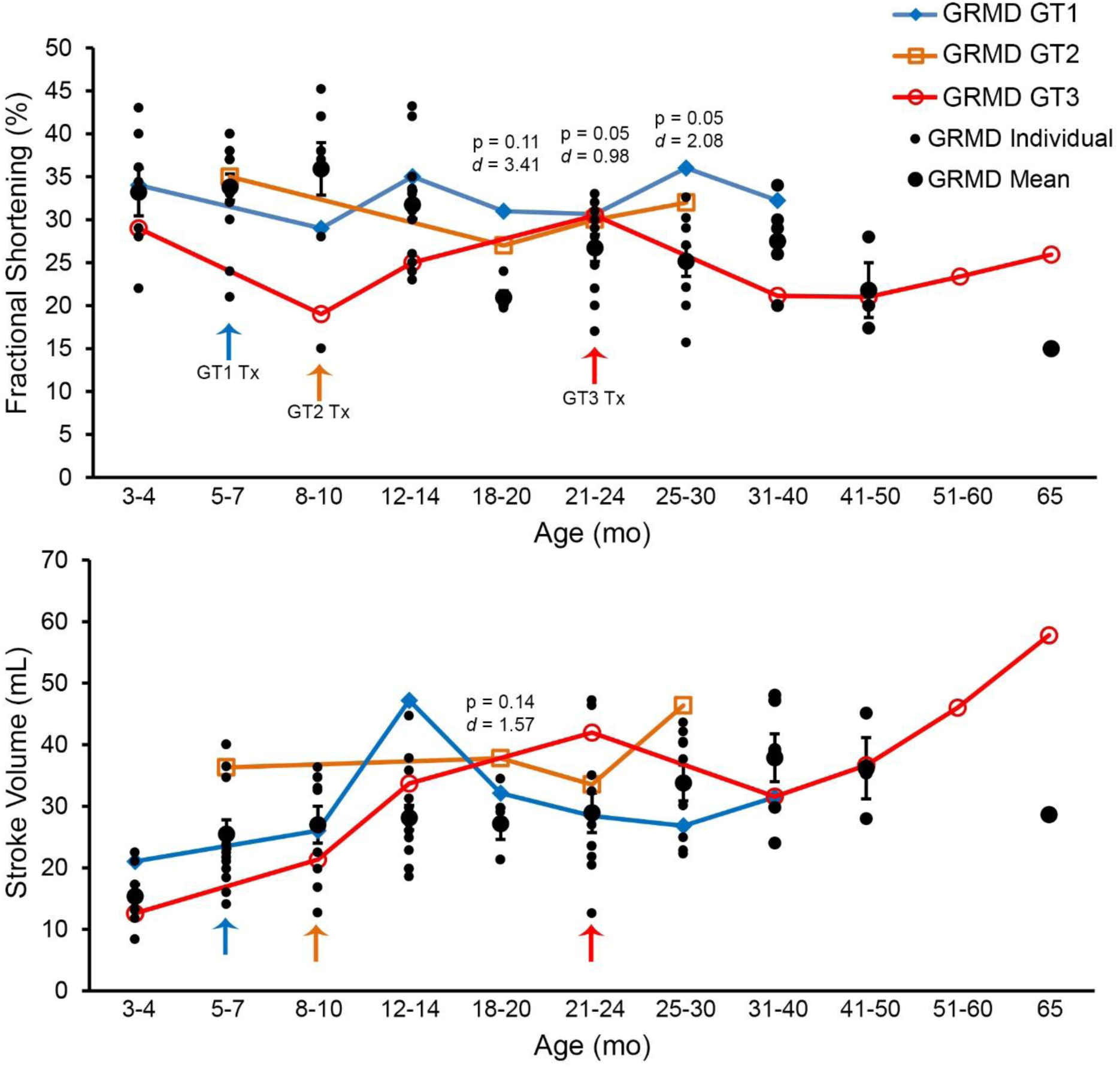
GRMD cardiac function following S100A1-ARC gene therapy. Fractional shortening and stroke volume measurements, as determined by echocardiography, from S100A1-ARC gene therapy treated GRMD dogs (GT1 in blue, GT2 in orange, and GT3 in red; color-matched arrows indicate respective age of treatment). Historical GRMD values from the colony for each age range are shown in black (means as large circles with SEM shown; individual values as small circles). Age ranges having two or more GT values were analyzed using two-tailed Welch’s T-test (α = 0.05) with effect size indicated.

